# SENOMORPHIC EFFECT OF GENETIC AND CHEMICAL PARTIAL REPROGRAMMING

**DOI:** 10.64898/2026.01.13.699211

**Authors:** Víctor Núñez-Quintela, Jeremy Chantrel, Miguel Ángel Prados, Pablo Pedrosa, Italo Lorandi, Lucía Lorenzo-Rodríguez, Cheng Chen, Raquel Paredes, Alejandro Failde-Fiestras, Diego González-Pérez, Alex Miralles-Domínguez, Sabela Da Silva-Álvarez, Ramón Lobato-Busto, Miguel González-Barcia, Clara Alcón, Joan Montero, Patricia Marqués, Dafni Chondronasiou, Federico Pietrocola, Manuel Serrano, Marta Kovatcheva, Aurora Gómez-Durán, Han Li, Manuel Collado

**Author notes:** Equal contribution (Víctor Núñez-Quintela, Jeremy Chantrel, Miguel Ángel Prados).

## Abstract

Partial reprogramming has emerged as a promising strategy to ameliorate aging phenotypes, yet its cellular targets and mechanisms remain poorly defined. Cellular senescence is a central hallmark of aging and a plausible mediator of reprogramming-induced rejuvenation. Here we show that genetic and chemical partial reprogramming act directly on senescent cells without restoring proliferative capacity. OSKM expression or a reduced two-compound regimen, tranylcypromine and RepSox (2c), attenuates senescence-associated secretory activity, restores mitochondrial homeostasis and apoptotic priming, and improves functional and inflammatory parameters in aged mice, establishing senomorphic, identity-preserving reprogramming as a potentially safer aging intervention.

## INTRODUCTION

The enforced expression of the Yamanaka factors Oct4, Sox2, Klf4, and c-Myc (OSKM) reprogram fully differentiated adult somatic cells into a pluripotent state, representing one of the most striking examples of cellular plasticity^1^. Although cells from aged organisms retain the capacity to undergo reprogramming, they do so with reduced efficiency^2^, highlighting aging as a major barrier to reprogramming. Importantly, reprogramming resets multiple molecular and cellular hallmarks associated with aging^3–5^; however, sustained OSKM expression results in full dedifferentiation and teratoma formation^6^, limiting its translational potential.

Recent advances demonstrated that short, intermittent induction of Yamanaka factors can uncouple epigenetic rejuvenation from loss of cell identity and ameliorate aging phenotypes in progeroid and naturally aged mice^7–10^, establishing partial reprogramming as a promising aging intervention strategy. In parallel, chemical reprogramming protocols capable of inducing pluripotency have been developed^11,12^, and a subset of this cocktail, composed of tranylcypromine (TCP) and RepSox (together referred to as 2c), has been claimed to be capable of affording the rejuvenation effects of complete reprogramming without altering somatic cell identity^13–15^. Despite these advances, the cellular targets and mechanisms through which partial reprogramming exerts its rejuvenating effects in vivo remain largely unknown^16,17^).

The accumulation of senescent cells is a hallmark of aging and a major driver of tissue dysfunction^18^, and their specific elimination has been shown to delay the onset and improve many aging-dependent pathologies^19,20^. Senescence is also intimately and bidirectionally linked to cellular reprogramming, acting both as an intrinsic barrier and as a potent extrinsic modulator of cellular plasticity^2,10,21–24^, raising the possibility that senescent cells may represent a key cellular target of reprogramming-based interventions. However, whether and how partial reprogramming directly remodels senescence phenotypes has not been systematically addressed.

Here, we directly tested whether partial reprogramming acts on the senescent state itself. Using enforced OSKM expression and a reduced chemical reprogramming strategy based on TCP and RepSox (2c), we show that partial reprogramming remodels senescent cells induced by different stimuli without restoring proliferative capacity. Both genetic and chemical approaches attenuate the abundance and functional activity of the senescence-associated secretory phenotype, restore mitochondrial homeostasis and apoptotic priming, and improve functional and inflammatory parameters in aged mice, while preserving senescence-associated cell-cycle arrest. Together, these findings establish partial reprogramming as a senomorphic intervention that decouples deleterious features of senescence from irreversible growth arrest. Moreover, we provide in vivo evidence that partial chemical reprogramming can improve selected aging phenotypes, supporting identity-preserving reprogramming as a potentially safer alternative to genetic approaches.

## RESULTS

### OSKM and 2c partially revert senescence transcriptome but not proliferation

To explore how cellular reprogramming impacts the cellular senescence phenotype, we forced the expression of the Yamanaka factors (Oct4, Sox2, Klf4 and c-Myc; OSKM) in mouse embryo fibroblasts (MEFs) previously induced to senescence by different stimuli: γ-radiation, nutlin-3 treatment, or HrasV12 expression (for oncogene-induced senescence, OIS) (Fig. 1a). OSKM expression was achieved either by lentiviral transduction of a doxycycline-inducible polycistronic OSKM cassette in wild-type MEFs^25^ or by doxycycline induction of a previously described transgenic OSKM cassette in MEFs derived from i4F mice^6^. Doxycycline was added to the cell culture three times for 48 h each, and cells were assessed at the end of the six-day treatment. Alternatively, we assessed chemical partial reprogramming by adding tranylcypromine (TCP) and RepSox (hereafter referred to as 2c) three times for a similar period of 6 days after which cells were collected and assessed.

**Fig 1.**
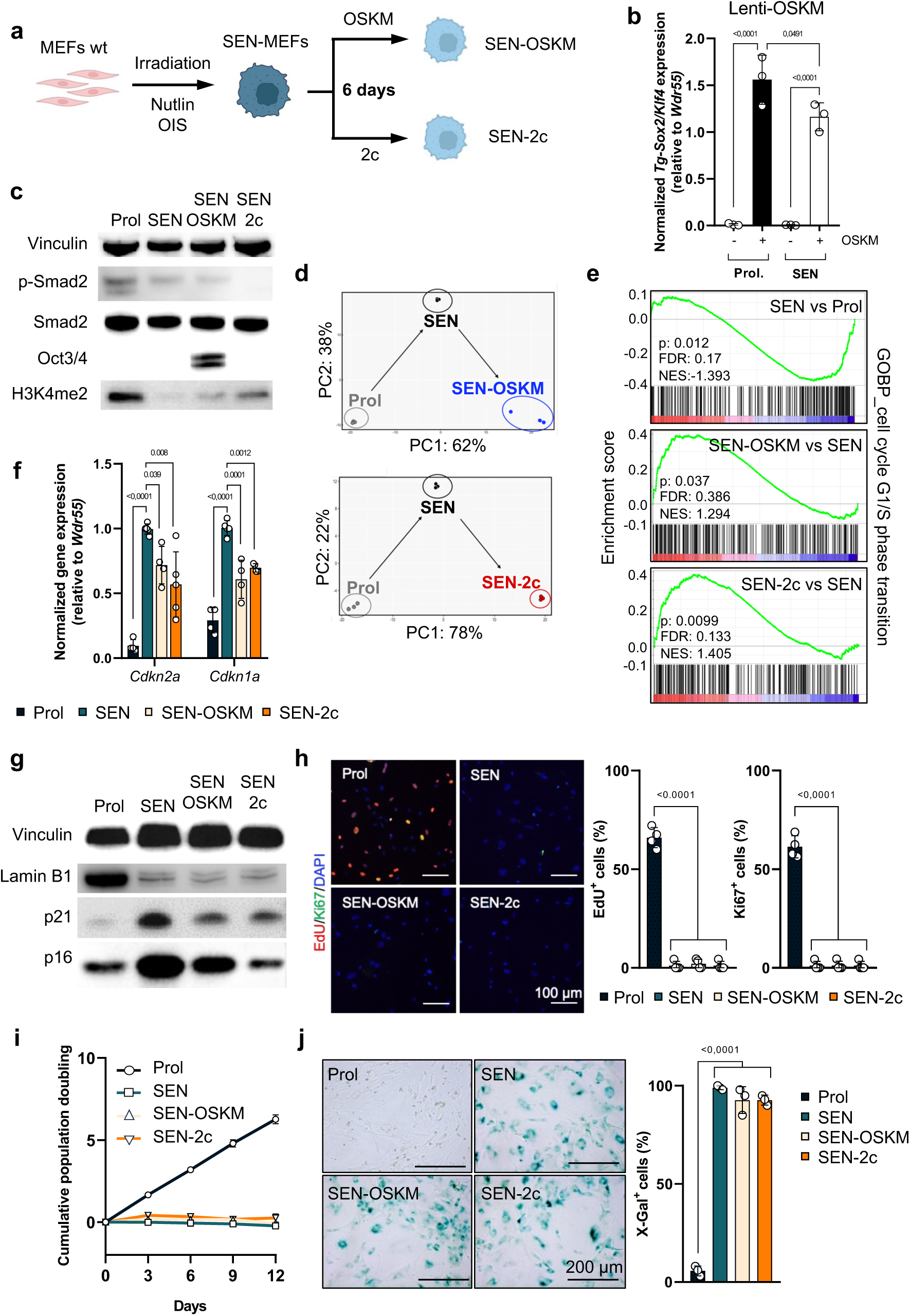
OSKM expression or 2c treatment partially reverts the senescence transcriptome without restoring proliferation. **a**, Schematic representation of the experimental design. **b**, RT–qPCR analysis of OSKM cassette expression (2^ΔΔCt^) in MEFs transduced with rtTA alone (-OSKM) or rtTA/OSKM (+OSKM). **c**, Immunoblot analysis of the indicated proteins across experimental conditions. Vinculin was used as a loading control. **d**, Principal component analysis of bulk RNA-seq data from proliferating and IR-induced senescent MEFs subjected to OSKM expression (top) or 2c treatment (bottom). **e**, Gene set enrichment analysis of the “Cell cycle G1/S phase transition” gene set. **f**, RT–qPCR analysis of *Cdkn2a* and *Cdkn1a* mRNA levels (2^ΔΔCt^), normalized to *Wdr55.* **g**, Immunoblot analysis of Lamin B1, p21, p16 across experimental conditions. Vinculin was used as a loading control. **h**, Representative immunofluorescence images of EdU incorporation and Ki67 staining, with corresponding quantification. Nuclei were counterstained with DAPI. Scale bar, 100 μm. **i**, Cumulative population doubling curves of MEFs across experimental conditions over the indicated time course. **j**, Senescence-associated β-galactosidase activity assessed by X-gal staining, with quantification. Scale bar, 200 μm. (**b, f, g, h** and **j**), Statistics were performed using one-way ANOVA followed by Tukey’s post hoc test. Error bars represent the s.d. For all experiments using MEFs, data represent at least three independent embryo-derived MEF lines with a minimum of two technical replicates per biological replicate.

First, we checked that the reprogramming cassette was being expressed by measuring its expression by RT-qPCR using oligonucleotides that are specific for mRNAs transcribed from the artificial cassette (Fig. 1b and Extended Data Fig. 1a). Consistently, we verified by Western blot that Oct4 protein was being expressed (Fig. 1c). Also, we measured the protein levels of P-SMAD2/SMAD2, as a readout of RepSox inhibition of TGFβR1, and of H3K4me2, indicative of LSD1 inhibition by TCP (Fig. 1c). The observed reduction in P-SMAD2 and the increase in H3K4me2 indicate that both, RepSox and TCP, were working as expected.

In order to have an unbiased view of the effect of OSKM expression or 2c treatment on the senescence phenotype, we performed bulk RNA-seq transcriptomics comparing proliferative MEFs, senescent controls, and senescent cells expressing OSKM or treated with 2c. Principal component analysis revealed that irradiation-induced senescent cells expressing OSKM or treated with 2c shifted away from untreated senescent controls and toward proliferating cells, although they remained clearly distinct from the proliferative state (Fig. 1d). Gene set enrichment analysis showed how several signatures closely linked to the senescent phenotype were partially reverted, such as “Cell cycle G1/S phase transition” (Fig. 1e) or “Mitotic DNA Integrity checkpoint signaling” (Extended Data Fig. 1b). Similarly, OSKM expression or 2c treatment caused a profound alteration in the transcriptional profile of OIS and nutlin-3-induced senescent cells (Extended Data Fig. 1c). We then decided to check the mRNA levels of cell cycle inhibitors *Cdkn2a* and *Cdkn1a* (coding for p16 and p21, respectively), the prototypical cell cycle inhibitors most commonly involved in implementing the proliferative arrest characteristic of cell senescence. RT-qPCR showed that mRNAs for both genes were partially, but consistently, reduced when OSKM was expressed on senescent cells or after 2c treatment (Fig. 1f). This effect was also corroborated at the protein level by analyzing p16 and p21 levels by Western blot (Fig. 1g). Interestingly, the levels of Lamin B1, a protein whose levels are frequently found downregulated during senescence, were not restored after OSKM expression or 2c treatment (Fig. 1g).

Although these transcriptional changes suggest a potential reversion, at least partial, of proliferative arrest, a defining feature of cellular senescence^18^, functional assays did not reveal restoration of proliferative capacity after either OSKM expression or 2c treatment. DNA synthesis assessed by EdU incorporation and immunofluorescence staining for the proliferation marker Ki67 remained undetectable in senescent cells expressing OSKM or 2c treatment (Fig. 1h and Extended Data Fig.1d). Furthermore, long-term growth curves of these senescent cells over 12 days in culture after OSKM expression or 2c treatment did not show any signs of resumed proliferation (Fig. 1i). Next, we determined whether OSKM expression or 2c treatment affected the senescence-associated beta-galactosidase activity (SAβGal), the most widely used marker of cell senescence^26^, using two strategies: a chromogenic substrate, X-gal, measured by cell staining; and a fluorescent probe, C_12_FDG, quantified by flow cytometry. In both cases, cells remained strongly positive for both substrates, indicative of unaltered SAβGal activity in irradiation-induced, OIS or nutlin-induced senescent cells (Fig. 1j and Extended Data Fig.1e,f).

Therefore, both OSKM-expression and 2c treatment induce a partial reversion of senescence-associated transcriptional programs. However, this transcriptomic alteration is insufficient to overcome two crucial characteristics of senescence: the stable proliferation arrest and sustained SAβGal activity.

### OSKM and 2c induce changes in the senescence phenotype and pattern of expression

Despite the persistent cell-cycle arrest and SAβGal positivity upon OSKM expression or 2c treatment, we realized that the morphology of the senescent cells was altered by both strategies.

We measured the cell size, typically enlarged in senescent cells^26^, and found it to be partially reduced when OSKM was expressed or when cells were treated with 2c (Fig. 2a). Similarly, the distinctive large nuclear area of senescent cells^27^ was almost completely restored to a normal, proliferating cell size, by OSKM or 2c (Fig. 2b). This led us to further analyze the expression of senescence markers, particularly various genes belonging to the senescence-associated secretory phenotype (SASP) and found that mRNAs for *Il6*, *Ccl2* and *Timp1*, were all partially reduced in a consistent manner by OSKM expression and 2c treatment (Fig. 2c). Moreover, using a cytokine array containing antibodies recognizing soluble secreted factors, and among them many members of the SASP, we identified as increased several of them (Serpine E1, Mmp3, Ccl5, etc.), as expected, when the membranes were incubated with conditioned media (CM) from senescent control cells compared to CM from proliferating ones. These same factors were found decreased in the CMs obtained from senescent cells expressing OSKM or treated with 2c (Fig. 2d). Further supporting the effect of partial reprogramming by OSKM expression or 2c treatment on the pro-inflammatory phenotype typical of senescence at the transcriptomic level, RNA-seq analysis revealed an attenuation of the SAUL_SEN_MAYO signature^28^, composed mainly of genes belonging to secreted pro-inflammatory factors (Fig. 2e and Extended Data Fig. 2a).

**Fig 2.**
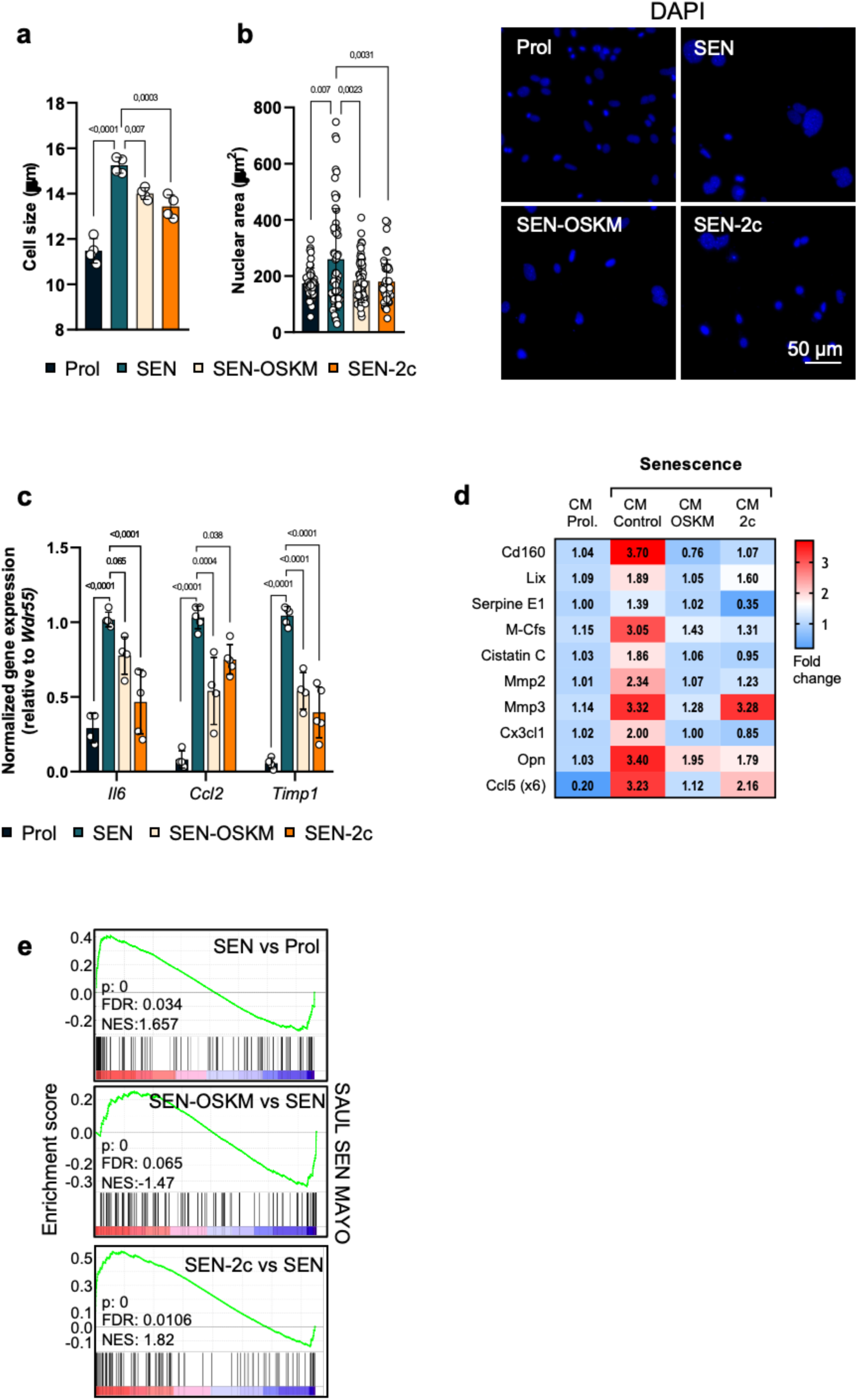
Partial reprogramming modulates senescence phenotypes. **a,** Quantification of cell area and **b**, nuclear area and representative immunofluorescent images of DAPI stained nuclei across the indicated conditions. **c,** RT–qPCR analysis of *Il6*, *Ccl2*, and *Timp1* mRNA levels (2^ΔΔCt^), normalized to *Wdr55*. **d,** Cytokine array analysis of conditioned media (CM) showing relative abundance of secreted SASP factors across the indicated conditions. **e,** Gene set enrichment analysis of the “Saul SenMayo” gene signature from bulk RNA-seq data. (**a–c**) Statistics were performed using one-way ANOVA followed by Tukey’s post hoc test. Error bars represent the s.d.

This reduction in the SASP composition after genetic of chemical partial reprogramming is not exclusive of MEFs since similar effects could be observed when whole bone marrow cells from mice, induced to senescence by doxorubicin treatment were forced to express OSK or were treated with 2c (Extended Data Fig. 2b). RT-QPCR showed that mRNA levels for SASP factors *Il6*, *Il1b* and *Tnfa* were all upregulated by doxorubicin treatment of bone marrow cells and both, OSK and 2c caused a reduction back to levels of normal, proliferating cells (Extended Data Fig. 2c). Moreover, similar effects were observed in human dermal fibroblasts expressing, in an inducible manner, progerin^29^, a pathological *LMNA* mutant frequently found in the progeric syndrome Hutchinson Gilford Progeria Syndrome: both OSK expression and 2c treatment led to a clear reduction of several SASP factors (Extended Data Fig. 2d,e).

Altogether, our results point to a partial reversion of the senescence phenotype caused by OSKM expression and a similar effect observed after 2c treatment, with a particular emphasis on the expression of secreted factors that conform the SASP.

### SASP activity is reduced by OSKM and 2c

Given the observed reduction in SASP components at the transcript and protein levels, we next wondered whether this reduction would translate into functional consequences. We collected CMs from control proliferative and senescent cells, and from senescent cells after expression of OSKM or 2c treatment, and used three well-established assays mediated by SASP factors: induction of paracrine senescence^30^, promotion of macrophage migration^31^, and enhancement of cellular reprogramming^22–24,32^ (Figure 3a).

**Fig 3.**
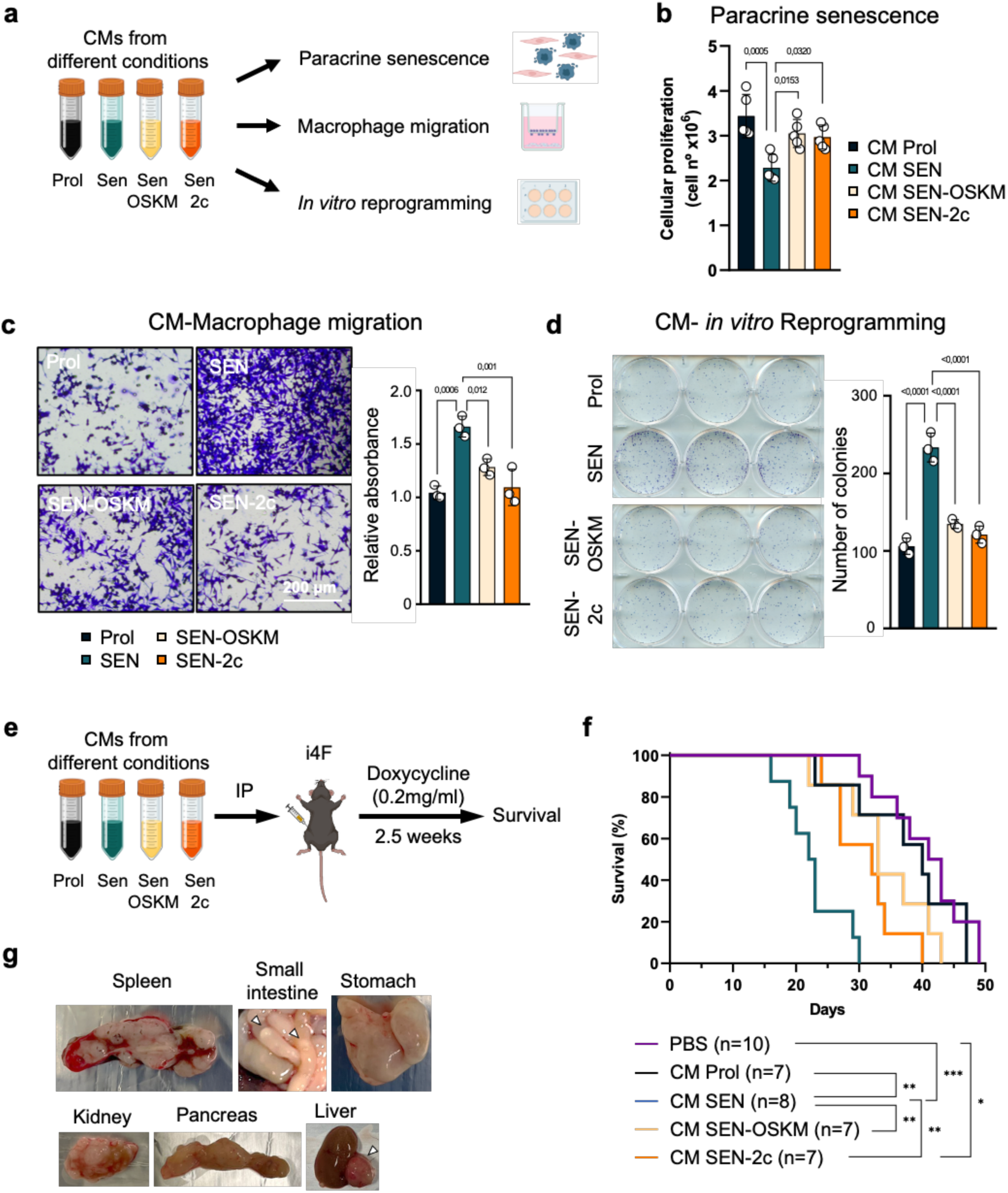
Partial reprogramming attenuates SASP functional activities. **a,** Schematic representation of the experimental designs for paracrine senescence, macrophage migration, and *in vitro* reprogramming assays using conditioned media (CM). **b,** Cellular proliferation of primary MEFs cultured with the indicated CM. **c,** Representative images (left) and quantification of macrophage migration (right) using RAW 264.7. Scale bar, 200 μm. **d,** Representative images of iPSC colonies (left) and quantification of colony numbers (right) in i4F MEFs cultured with the indicated CM. **e,** Schematic representation of the *in vivo* reprogramming assay in i4F mice. **f,** Kaplan-Meier survival curves of i4F mice. Sample sizes (*n*) for each group are indicated in the panel. **g,** Representative macroscopic images of teratomas from different organs of i4F mice; arrowheads indicate teratoma locations. (**b–d**) Statistics were performed using one-way ANOVA followed by Tukey’s post hoc test. Error bars represent the s.d. (**f**) Statistical significance was determined using the log-rank (Mantel–Cox) test. Statistical significance: *P < 0.05, **P < 0.01, ***P < 0.001.

To assess paracrine senescence, we applied these CMs to early-passage primary MEFs cultures, and paracrine senescence induction was evaluated. First, we measured cell proliferation as a surrogate marker of the induction of paracrine senescence and observed that CM from senescent control cells led to a marked reduction in proliferation of recipient MEFs compared to CM from proliferating cells. In contrast, CM derived from senescent cells expressing OSKM or treated with 2c caused a significantly weaker growth-inhibitory effect (Fig. 3b). We also measured SAβGal activity on the recipient cells by flow cytometry using the fluorescent substrate C12FDG. MEFs receiving CM from senescent control cells showed a marked increase in SAβGal activity compared to those receiving CM from control proliferative cells, as expected (Extended Data Fig. 3a,b). However, when CMs from senescent cells expressing OSKM or treated with 2c were added, recipient MEFs showed clearly a lower level of SAβGal activity, indicative of a reduced induction of paracrine senescence (Extended Data Fig. 3a,b). To further substantiate this result, we measured the cell size of cells receiving these CMs. Cells cultured with the senescent control CM showed a larger size than those receiving control proliferative CM, as expected for cells undergoing paracrine senescence (Extended Data Fig. 3c). However, when the CM came from senescent cells expressing OSKM or treated with 2c, the size of the recipient cells remained similar to the ones receiving control proliferative CM (Extended Data Fig. 3c). This attenuation of the paracrine senescence activity of the SASP by OSKM expression or 2c treatment was also evidenced at the molecular level by the reduced expression of mRNAs coding for cell cycle inhibitors *Cdkn2a* and *Cdkn1a*, and SASP factors *Il6*, *Ccl2* and *Areg* (Extended Data Fig. 3d) and the reduced presence of the DNA damage marker γH2AX (Extended Data Fig. 3e).

Another documented effect of the SASP is promoting macrophage migration^31^. Using transwell migration assays (Fig. 3a), we confirmed that RAW264.7 macrophages migrated more efficiently when control irradiation-induced senescent MEFs were on the bottom chamber compared to the presence of control proliferative ones (Fig. 3c). When we placed senescent MEFs expressing OSKM or previously treated with 2c in the bottom chamber, the migration of RAW 264.7 macrophages was similar to the one observed with control proliferative cells (Fig. 3c). Similar results were also obtained when we used OIS or nutlin-induced senescence CM added to bone marrow derived macrophages (Extended Data Fig. 3f).

Although cell senescence is an intrinsic barrier for reprogramming, senescent cells can enhance reprogramming of neighboring cells through secreted SASP factors^22–24,32^. Therefore, we assessed whether SASP released from senescent cells after OSKM expression or 2c treatment retains this reprogramming-enhancing activity (Fig. 3a). MEFs derived from i4F mice were cultured in CMs from control proliferative and senescent cells, or senescent cells expressing OSKM or treated with 2c, followed by doxycycline-induced activation of the OSKM transgenic cassette. As previously described, reprogrammable MEFs receiving CM from senescent control cells showed a clear enhancement of the reprogramming efficiency compared to the ones cultured with CM from the proliferative control as evidenced by the higher number of iPSC colonies obtained. Consistently, the CMs from senescent cells expressing OSKM or treated with 2c lost this capacity to enhance reprogramming (Fig. 3d and Extended Data Fig. 3g).

We extended these findings in vivo by intraperitoneally injecting CM into i4F reprogrammable mice, followed by doxycycline administration in the drinking water to induce OSKM expression and teratoma formation as a readout of in vivo reprogramming (Fig. 3e). Animals receiving CM from control senescent cells developed teratomas much faster and succumbed to tumors earlier than PBS-injected mice or those receiving CM from control proliferative MEFs (Fig. 3f,g). In contrast, CM from senescent cells expressing OSKM or treated with 2c significantly delayed teratoma formation and prolonged survival (Fig. 3f), implying the loss of SASP-mediated enhancement of reprogramming activity.

All these in vitro and in vivo systems concurrently prove that senescent cells expressing OSKM or treated with 2c not only show reduced abundance of SASP factors secretion but also present a weaker SASP functional activity, as measured by paracrine senescence induction, promotion of macrophage migration and enhancement of cell reprogramming.

### OSKM and 2c revert the mitochondrial phenotype of senescence

Recent work has demonstrated that mitochondrial dysfunction is a hallmark of senescence and a key driver of the SASP^33–37^. The alteration of mitochondria in senescence is characterized by an increase in mitochondrial mass that is considered an attempt by senescent cells to compensate for the loss of mitochondrial function^38,39^. This is measured by flow cytometry using mitotracker, a probe that reflects mitochondrial mass, combined with TMRM, a probe detecting mitochondrial membrane potential and indicative of function. Consistently, we found that senescent cells showed higher mitotracker fluorescent signal compared to proliferative cells (Fig. 4a). In contrast, senescent cells expressing OSKM or treated with 2c presented a mitochondrial mass similar to the one observed in proliferative control cells (Fig. 4a). In addition, senescent control cells showed higher levels of TMRM that could derive from their increased mitochondrial mass (Extended Data Fig. 4a). Since this could be concealing the reduction in mitochondrial membrane potential, we analyzed the TMRM to mitotracker ratio. This allowed us to clearly observe the mitochondrial dysfunction of senescent control cells and how this was reverted by OSKM expression or 2c treatment (Fig. 4b).

**Fig 4.**
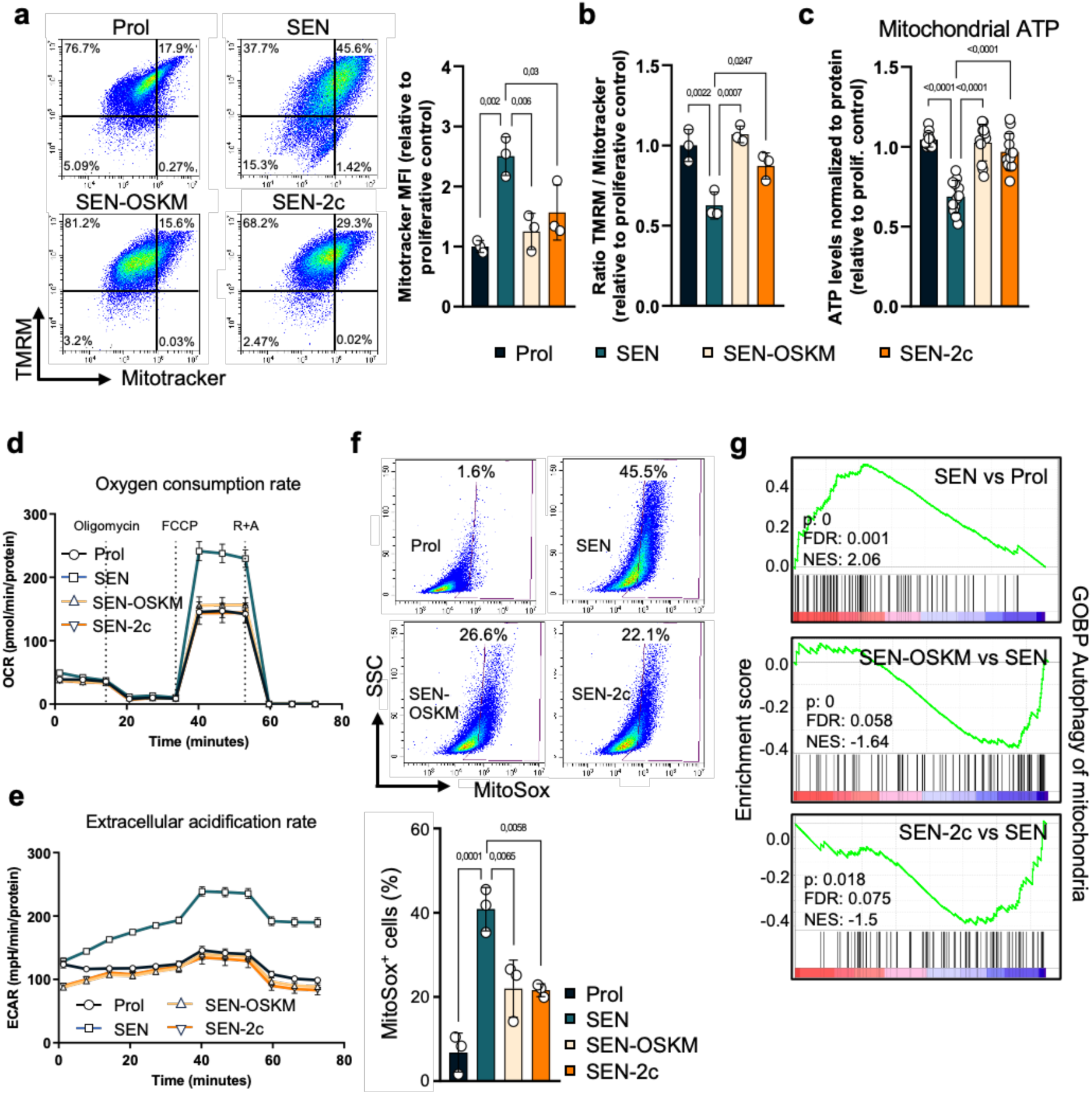
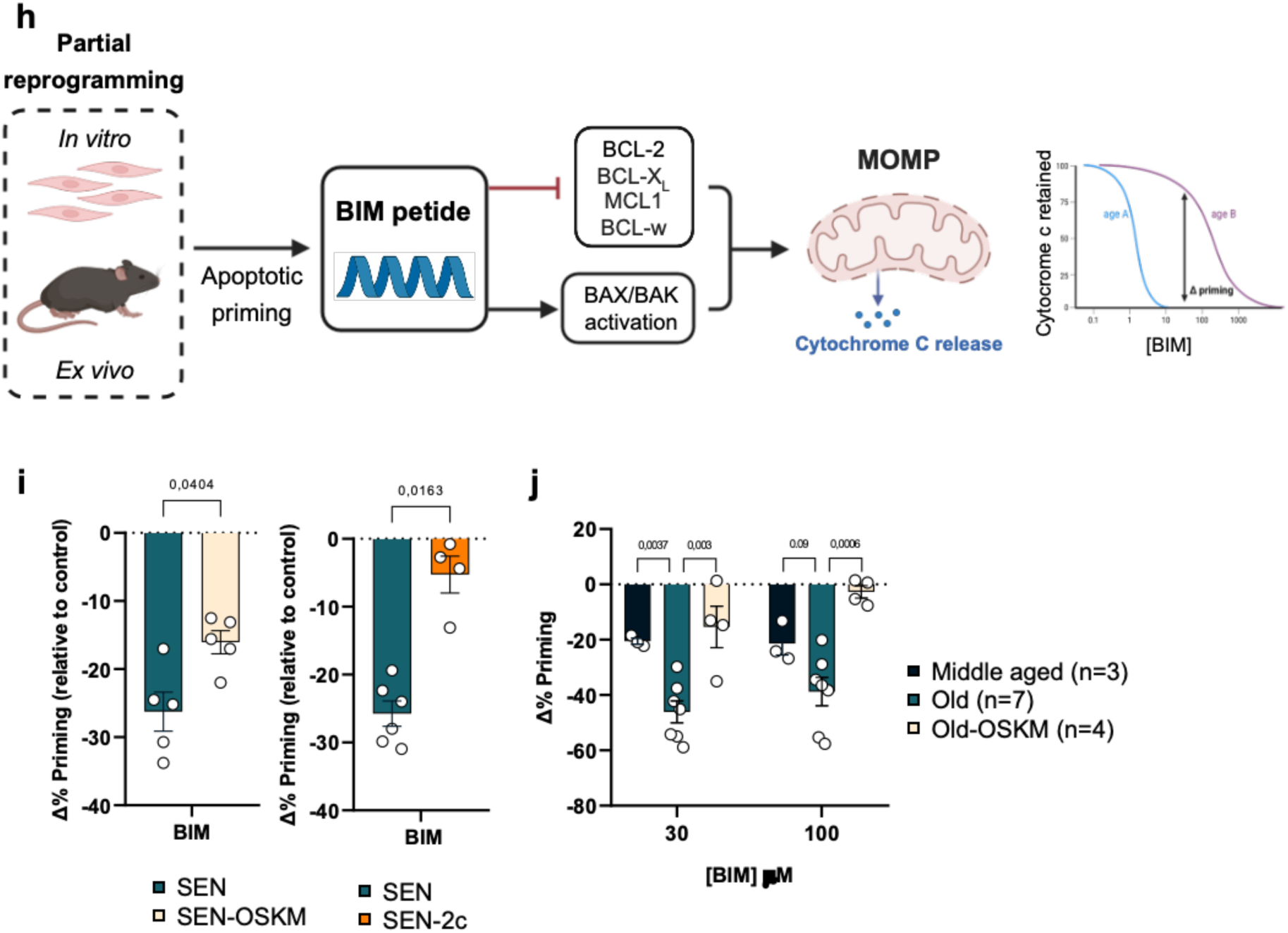
Partial reprogramming restores mitochondrial homeostasis and apoptotic sensitivity. **a,** Representative flow cytometry plots (left) and quantification of Mitotracker Mean Fluorescence Intensity (MFI) (right) across experimental conditions. **b,** Ratio of TMRM to Mitotracker fluorescence. **c,** Mitochondrial ATP levels normalized to total protein content. **d,** Oxygen consumption rate measurements. **e**, Extracellular acidification rate measurements. **f,** Representative flow cytometry plots (top) and quantification (bottom) of MitoSox^+^ cells. **g,** Gene set enrichment analysis of the “GOBP Autophagy of mitochondria” gene set. **h,** Schematic representation of the apoptotic priming assessment. **i**, Apoptotic priming (Δ priming) measured by BH3 profiling in MEFs in response to BIM peptide. **j,** Δ priming in adipocytes isolated from visceral white adipose tissue of mice (*n*=3 middle aged, *n*=7 old, *n*=4 old-OSKM). (**a–c, f, i** and **j**) Statistics were performed using one-way ANOVA followed by Tukey’s post hoc test. Error bars represent the s.d.

We next decided to further characterize other mitochondrial parameters. When analyzing mitochondrial ATP levels, we observed a decrease in control senescent cells compared to the proliferative condition, whereas in senescent cells expressing OSKM or treated with 2c, ATP levels were restored to those showed by the proliferative control (Fig. 4c). The analysis of oxygen consumption under these conditions, revealed differences in maximal oxygen consumption rate only in control senescent cells after uncoupling the electron transport chain with FCCP (Fig. 4d). This implies that this increase in maximal respiratory capacity did not correlate with enhanced energy production in senescent cells. Altogether, these data demonstrate mitochondrial dysfunction during senescence, with a direct impact on energy production through oxidative phosphorylation, which is rescued by OSKM expression or 2c treatment.

Subsequently, we evaluated the extracellular acidification rate, a parameter directly related to glycolysis^40^. These data show that senescent cells display a clear increase in extracellular medium acidification compared to the proliferative condition, indicative of an upregulation of glycolysis during senescence (Fig. 4e). Again, this metabolic shift observed during senescence was reversed upon OSKM expression or 2c treatment (Fig. 4e).

Mitochondrial dysfunction in senescent cells is also associated with increased production of reactive oxygen species (ROS), which contributes to the oxidative damage suffered by many key cellular components that is characteristic of cellular senescence^38,41^. Using MitoSox, a fluorescent probe that detects superoxide, we observed elevated levels of mitochondrial ROS in control senescent cells compared to proliferative ones, as expected, whereas expression of OSKM or 2c treatment resulted in reduced levels of MitoSox (Fig. 4f). Supporting these findings, transcriptomic analysis revealed dysregulation of mitophagy-related gene signatures during senescence, which were restored toward a proliferative-like state following OSKM expression or 2c treatment (Fig. 4g).

Resistance to apoptosis is another hallmark of senescent cells^18^. This resistance is frequently driven by transcriptional upregulation of mitochondrial anti-apoptotic genes^42,43^. Given the restoration of mitochondrial function by OSKM expression or 2c treatment, we next asked whether apoptotic sensitivity could also be reinstated in senescent cells. To assess apoptotic priming, we employed BH3 profiling, a single-cell, flow cytometry-based functional assay that quantifies mitochondrial outer membrane permeabilization (MOMP)^44^ in response to pro-apoptotic BH3 peptides, using cytochrome c release as a readout for apoptosis induction^45,46^. We used the pan-activating BH3 peptide Bim, which inhibits all major anti-apoptotic BCL-2 family members (Bcl-2, Bcl-xL, Mcl-1, and Bcl-w) and directly activates the pro-apoptotic effectors Bax and Bak, thereby inducing MOMP (Fig. 4h). We have recently shown that BH3 profiling can be applied to determine cell-type specific anti-apoptotic adaptations in senescent melanoma cell lines^47^.

Irradiation-induced senescent MEFs displayed resistance to apoptosis following exposure to the pan-activating BH3 peptide Bim, as evidenced by increased cytochrome c retention and a negative Δ% priming compared to proliferating cells (Fig. 4i and Extended Data Fig. 4b). Strikingly, senescent cells expressing OSKM or treated with 2c exhibited apoptotic priming profiles comparable to proliferating controls, with reduced cytochrome c retention and a less negative Δ% priming (Fig. 4i and Extended Data Fig. 4b). These results indicate that OSKM expression or 2c treatment restores apoptosis sensitivity in senescent cells.

We next asked whether similar changes in apoptotic priming occur during natural aging in vivo and whether they are reversible by reprogramming. While changes in BH3 profiling have previously been reported in various tissues during murine development^48^, this has never been investigated in aged mice. We performed BH3 profiling on the visceral white adipose tissue from young (12 weeks) and aged (90-114 weeks) mice, a tissue known to accumulate senescent cells during aging^20^. Adipocytes isolated from aged mice showed a negative Δ% apoptotic priming compared to young mice (Fig. 4j). Remarkably, these changes in apoptotic priming were already detectable in middle-aged mice (25 weeks), suggesting early acquisition of mitochondrial anti-apoptotic adaptation in the visceral fat tissue. Importantly, resistance to apoptosis in the aged mice was restored after a single cycle of OSKM reprogramming protocol (Fig. 4j), which we have previously reported to induce molecular features of rejuvenation^49^. Together, these data demonstrate that BH3 profiling can detect senescence-associated changes in apoptotic priming during natural aging, and that these alterations are reversible through partial OSKM reprogramming.

Collectively, our data show that OSKM expression and 2c treatment impact the mitochondria of senescent cells restoring its metabolic function and apoptosis sensitivity, providing a mechanistic basis for the reduced SASP factor release and activity.

### Distinct transcriptional responses to OSKM and 2c in senescent cells

To define how OSKM expression and 2c treatment remodel the transcriptional landscape of senescent cells, we analyzed bulk RNA-seq datasets from proliferating controls, senescent cells, and senescent cells subjected to either intervention. Senescence-associated differentially expressed genes (DEGs) were first identified by comparing senescent cells to proliferating controls. These genes were then classified based on how each intervention altered the senescence-associated transcriptional changes.

Categorization was based on FDR-adjusted significance (≤ 0.05) and effect size (|log2FC| > 1) across senescent–proliferative and intervention–senescent contrasts, with full reversion defined by loss of significant difference relative to proliferating controls. Four categories were defined: (1) reverted, in which expression was significantly shifted in the opposite direction of the senescence-associated change and no longer differed from proliferating controls; (2) partially reverted, in which expression was shifted toward proliferative levels but remained significantly different; (3) non-reverted, showing no significant change relative to senescence; and (4) further changed, in which expression was significantly altered in the same direction as the senescence-associated change (Extended Data Fig. 5a,b). Of note, OSKM expression affected a substantially larger fraction of senescence-altered genes than 2c treatment, indicating broader transcriptional remodeling (Fig. 5a). Directional changes underlying DEG categorization were consistent across biological replicates.

**Fig 5.**
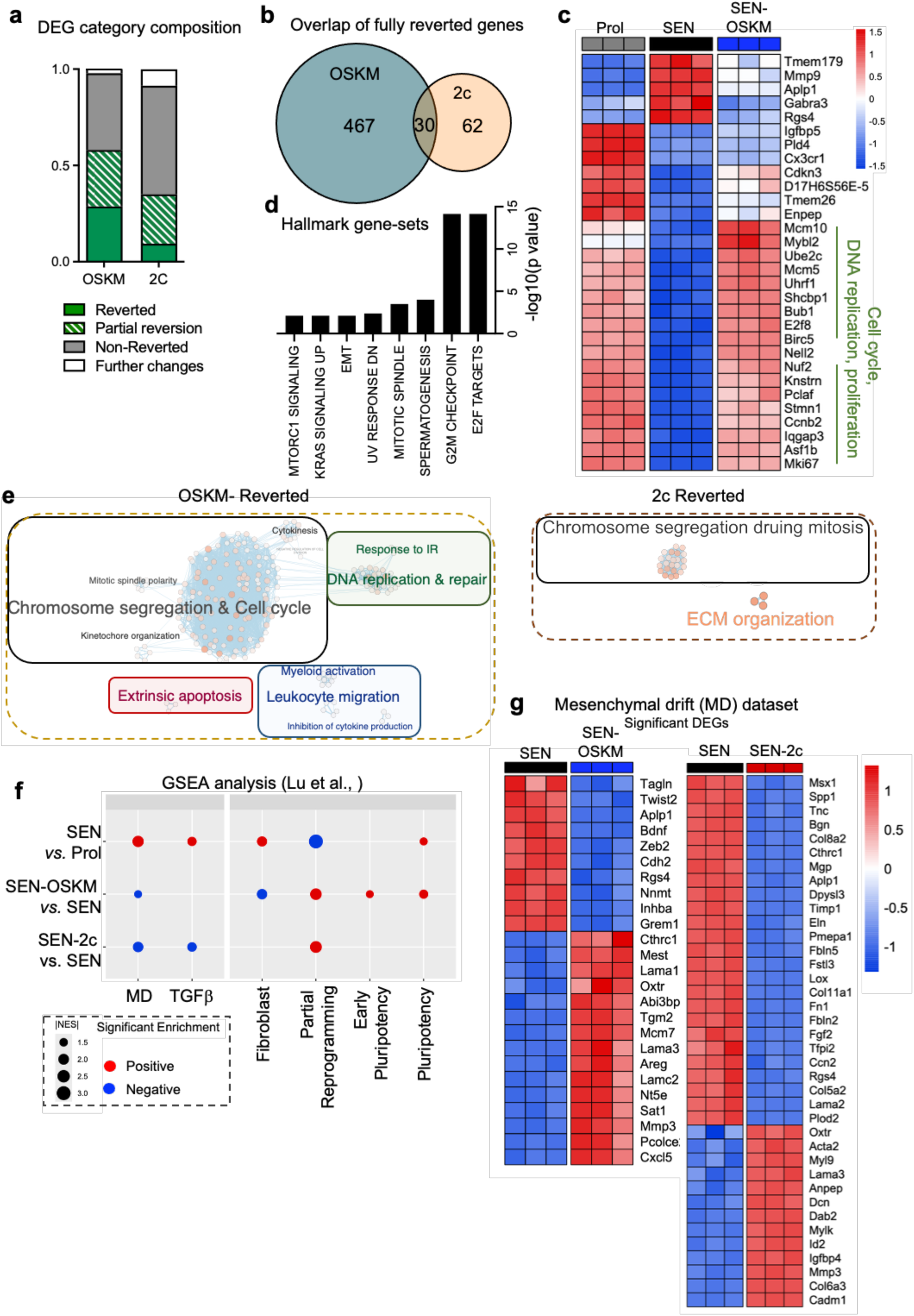
Partial reprogramming drives distinct transcriptional remodeling of the senescence landscape. **a,** Stacked bar plot showing the proportions of senescence-associated differentially expressed genes (DEGs) categorized by transcriptional response to OSKM expression or 2c treatment (reverted, partial reversion, non-reverted, or further changes). **b,** Venn diagram illustrating the overlap of fully reverted genes between OSKM and 2c interventions. **c,** Heatmap of relative expression for representative shared reverted genes involved in DNA replication and cell cycle progression (Z-score normalized). **d,** Functional enrichment analysis of the 30 shared reverted genes identified in (**b**). **e,** Bubble plots depicting biological pathways significantly reverted by OSKM expression (left) or 2c treatment (right). **f,** Dot plot summarizing Gene Set Enrichment Analysis results for mesenchymal drift (MD), TGFβ-related, and cell identity-related gene signatures. Dot size represents the absolute Normalized Enrichment Score (|NES|). **g,** Heatmaps showing the expression of significant MD-related DEGs in senescent cells compared to OSKM-treated (left) or 2c-treated (right) senescent cells.

Analysis of fully reverted genes revealed limited overlap between OSKM- and 2c-responsive sets (Fig. 5b,c). Gene set enrichment analysis (GSEA) of the 30 shared reverted genes showed enrichment for cell-cycle progression, DNA replication, and proliferative programs (Fig. 5d). Consistent with this, pathway-level analysis indicated that both interventions preferentially modulated cell-cycle–related pathways while also exhibiting intervention-specific effects. OSKM uniquely reverted pathways linked to apoptosis, DNA repair, and leukocyte migration, whereas 2c preferentially affected extracellular matrix organization (Fig. 5e, Supplementary Table 6).

### Selective suppression of mesenchymal drift by 2c without loss of somatic identity

Aging has been associated with chronic activation of mesenchymal and EMT-related transcriptional programs, termed mesenchymal drift (MD), which correlates with fibrotic disease severity and reduced survival^50^. In this framework, MD reflects sustained activation of EMT-derived transcriptional programs, quantified using the Hallmark EMT gene set supplemented with key EMT transcription factors. OSKM-mediated reprogramming has been shown to suppress MD scores prior to overt dedifferentiation and pluripotency acquisition^50^. Given that senescent cells frequently upregulate EMT- and TGFβ-driven transcriptional programs^51,52^, we examined whether senescence is associated with MD-related signatures and whether these are modulated by OSKM or 2c.

Applying MD- and TGFβ-related gene signatures from Lu et al., we observed that senescent cells exhibited significantly elevated MD and TGFβ scores relative to proliferative controls. Both OSKM expression and 2c treatment significantly reduced the senescence-associated MD score (Fig. 5f,g, Supplementary Table 6). In contrast, only 2c treatment significantly suppressed the TGFβ signature, whereas OSKM did not, suggesting that the two interventions attenuate mesenchymal-associated aging signatures through partially distinct transcriptional mechanisms.

We next assessed whether OSKM or 2c induced transcriptional states associated with partial reprogramming state as previously defined^50^. Both interventions increased partial reprogramming scores in senescent cells, consistent with engagement of early reprogramming-associated changes. OSKM expression additionally increased early and full pluripotency-associated signatures and suppressed fibroblast identity programs in senescent cells, whereas 2c treatment did not elicit comparable identity-related changes (Fig. 5f and Extended Data Fig. 5c,d). Thus, 2c selectively attenuates senescence-associated MD and TGFβ activation without inducing broader shifts in somatic identity.

Together, these results show that senescent cells display sustained activation of EMT-derived transcriptional programs consistent with MD shift and that both OSKM and 2c suppress this state, with 2c acting through a more selective transcriptional remodeling program.

### 2c improves motor performance and behavioral parameters in old mice

Based on our preceding results implicating 2c in the modulation of senescence- and aging-associated mechanisms, we investigated the effects of chemical partial reprogramming *in vivo.* To this end, 29-month-old HET3 female mice were treated with the 2c protocol, and a range of physiological, behavioral, and molecular parameters associated with aging were assessed (Fig. 6a).

**Fig 6.**
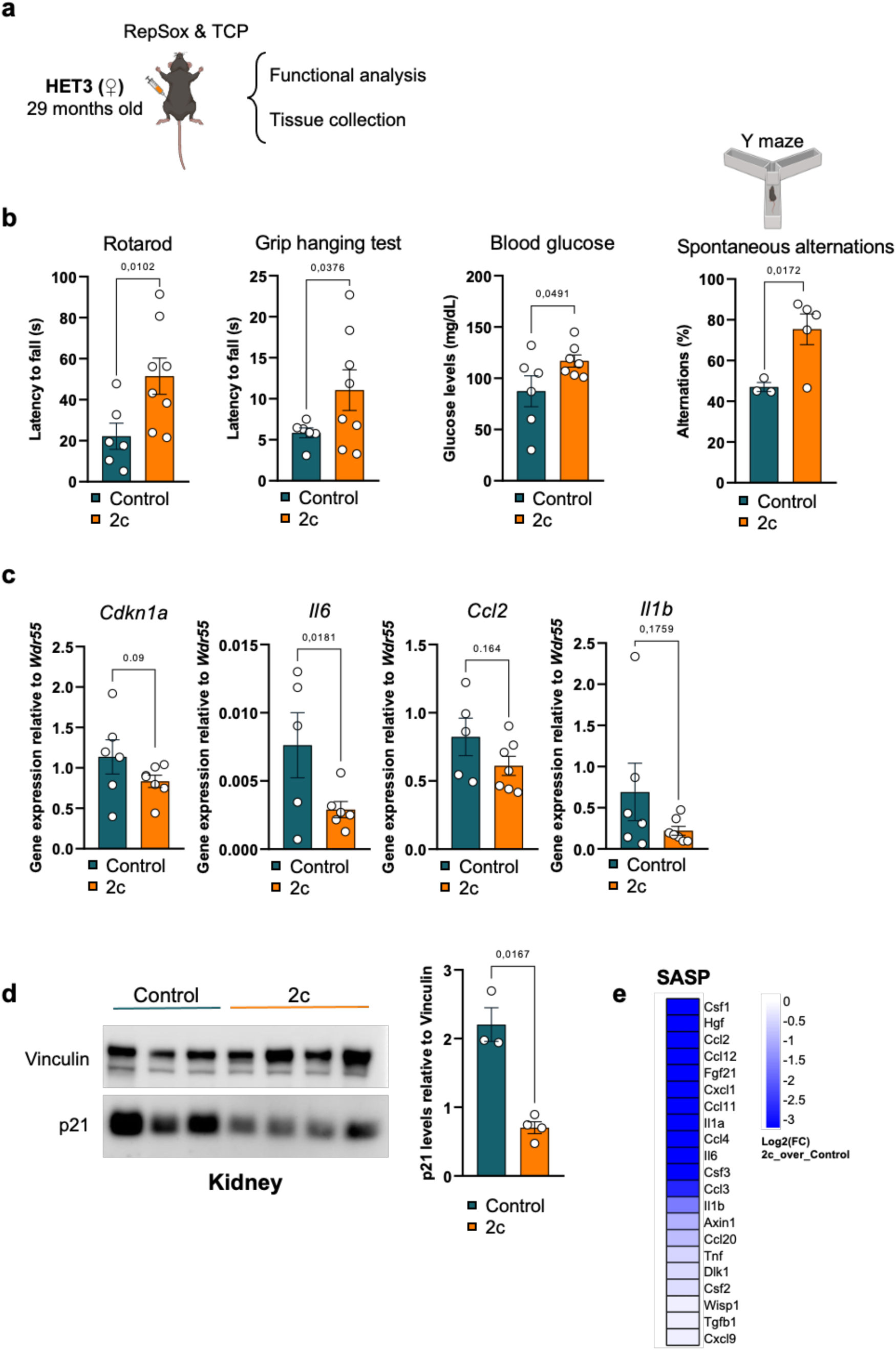
Chemical partial reprogramming improves physical function and attenuates aging biomarkers in old mice. **a,** Schematic representation of the experimental design for *in vivo* 2c treatment in aged (29-month-old) HET3 female mice. **b,** Functional and behavioral assessments of control and 2c-treated aged mice. **c,** RT–qPCR analysis of hepatic *Cdkn1a*, *Il6*, *Ccl2*, and *Il1b* mRNA levels (2^ΔΔCt^) normalized to *Wdr55*. **d,** Immunoblot analysis (left) and corresponding quantification (right) of p21 protein levels in kidney lysates. Vinculin was used as a loading control. **e,** Heatmap showing the relative abundance (log_2_ fold change) of serum SASP factors detected by Olink proteomic profiling. (**b–d**) Statistics were performed using unpaired two-tailed t-test. Error bars represent the s.e.m. Sample size: Control group, *n*=6; 2c group, *n*=8.

Treatment with 2c resulted in improved physical performance in aged mice. In the accelerating rotarod test, 2c-treated animals showed a significantly longer latency to fall than old mice controls, indicating better motor coordination and balance (Fig. 6b). Similarly, in the grip hanging test, 2c-treated mice exhibited an increased latency to fall, consistent with enhanced muscle strength (Fig. 6b). In parallel, 2c-treated animals exhibited a recovery of fasting glucose levels, suggesting an improvement in glucose homeostasis in old mice, although the underlying mechanism remains unclear (Fig. 6b). In the Y-maze test, 2c-treated mice exhibited a significantly higher percentage of spontaneous alternations compared to controls, consistent with an improvement in spatial working memory (Fig. 6b). Other tests included the open-field test, in which 2c-treated mice showed a clear tendency towards increased total distance travelled and higher maximum speed, although the number of entries into the central zone did not differ between groups suggesting that the increase in locomotor activity is not accompanied by major changes in anxiety-like behavior; and the elevated zero maze, with 2c-treated mice showing a tendency to travel longer distances and to spend more time in the open area, and with only mice in this group fully crossing the open segment (Extended Fig. 6a,b).

Consistent with the systemic functional improvements observed in 2c-treated mice, molecular analyses revealed reduced expression of senescence and inflammation-associated genes in several tissues. In the liver, 2c treatment led to a significant decrease in *Il6* transcript levels and a downward trend in *Cdkn1a*, *Ccl2, Il1b, Cxcl1 and Il1a* expression compared with aged controls (Fig. 6c and Extended Fig. 6c), while other classic markers of senescence, such as *Cdkn2a*, did not show any reduction in expression (Extended Fig. 6c). This attenuation of senescence markers was further supported by reduced p21 protein abundance in the kidney of 2c-treated animals (Fig. 6d). Moreover, proteomic profiling of serum revealed the reduction of circulating factors (Extended Fig. 6d), including key SASP pro-inflammatory cytokines and chemokines in mice receiving 2c (Fig. 6e), indicating that partial reprogramming with 2c mitigates molecular hallmarks of aging across multiple organs and dampens systemic inflammatory signaling in old mice. Interestingly, 2c does all this without causing any detectable health issue, as judged by a maintained body weight (Extended Fig. 6e), and without an apparent reduction of senescent cells, as judged by SAβGal staining of white adipose tissue (Extended Fig. 6f). Taken together, these data suggest that partial chemical reprogramming by 2c treatment mitigates molecular and functional hallmarks of aging across multiple organs and dampens systemic inflammatory signaling in old mice.

## DISCUSSION

Partial reprogramming by cyclic and transient expression of Yamanaka factors has emerged as a promising strategy to ameliorate aging-associated phenotypes^17^. Previous studies have shown that short-term OSKM expression can uncouple epigenetic rejuvenation from permanent dedifferentiation and improve molecular and functional features of aging in progeroid and naturally aged mice ^7–10^. However, critical questions remain regarding the safety of gene-based interventions, which aged cell types and states are responsive, and how core hallmarks of aging are remodeled at the cellular and tissue levels.

Given that cellular senescence is a central driver of aging and age-related diseases^18^ and is intimately linked to reprogramming^2,10,21–24^, we investigated whether partial reprogramming directly alters senescence phenotypes. By comparing genetic (OSKM) and chemical (TCP+RepSox; 2c) partial reprogramming strategies across multiple senescence cellular models and in vivo aging paradigms, we show that both interventions suppress inflammatory SASP abundance and activity, improve mitochondrial dysfunction, and partially reset apoptotic priming, while preserving stable growth arrest. These findings indicate that core senescence blocks persist despite broad molecular remodeling and support a senomorphic mode of action. Notably, 2c induces a more selective transcriptional remodeling, reducing mesenchymal drift and TGFβ signaling without erasing somatic identity, and improves functional and molecular aging-associated parameters in very old mice without apparent reduction of senescent-cell burden.

Consistent with a previous report^53^, partial reprogramming did not reverse two defining hallmarks of senescence: stable cell-cycle arrest and sustained SAβGal activity. The persistence of proliferative arrest despite partial normalization of cell-cycle–associated gene expression likely reflects the multilayered nature of senescence stability^18^. Senescent cells harbor persistent DNA damage responses, telomere-associated lesions, chromatin and nuclear lamina alterations, and reinforced checkpoint circuitry that may not be readily reversible by short-term reprogramming interventions. In line with this, Lamin B1 levels, a marker of senescence-associated nuclear remodeling, remained suppressed following OSKM expression or 2c treatment. Thus, while partial reprogramming engages transcriptional programs associated with proliferation, it does not overcome the structural and checkpoint barriers that safeguard against aberrant cell-cycle re-entry in stressed cells.

In contrast to the irreversibility of growth arrest, both genetic and chemical partial reprogramming exerted a pronounced and functionally relevant effect on the SASP program. OSKM expression and 2c treatment reduced the expression and secretion of multiple SASP components both in vitro and in vivo, leading to diminished SASP activity across several functional assays. Conditioned media from treated senescent cells showed reduced capacity to induce paracrine senescence, promote macrophage migration, and enhance cellular reprogramming efficiency in neighboring cells, indicating that partial reprogramming suppresses not only SASP abundance but also its biological impact.

Mechanistically, our data are consistent with a model in which restoration of mitochondrial homeostasis contributes to SASP attenuation. Both OSKM and 2c partially normalized various senescence-associated mitochondrial dysfunctions^33–37^, including impaired oxidative phosphorylation, elevated mitochondrial ROS, and altered metabolic flux. While SASP regulation is multifactorial^18^, mitochondrial remodeling provides a plausible upstream mechanism linking partial reprogramming to reduced inflammatory output. The mitochondrial remodeling induced by 2c may, at least in part, reflect direct effects of its constituent compounds. Tranylcypromine (TCP) is a nonselective, irreversible inhibitor of monoamine oxidases (MAOs)^54^, which reside on the outer mitochondrial membrane and have been implicated in mitochondrial dysfunction and senescence induction^55^. Specifically, MAO-A upregulation has been shown to promote mitochondrial oxidative stress and senescence in cardiomyocytes^55^. Although TCP exhibits a modest preference for MAO-B, inhibition of MAO activity could plausibly contribute to the improved mitochondrial function observed in senescent cells following 2c treatment. Disentangling these pharmacological effects from reprogramming-associated transcriptional remodeling will be important for defining the mechanisms underlying chemical partial reprogramming.

Mitochondrial remodeling was accompanied by reversal of senescence-associated apoptotic resistance^18^, as evidenced by restored apoptotic priming in vitro and in adipocytes from aged mice following partial OSKM reprogramming. While BH3 profiling measures mitochondrial readiness rather than execution of apoptosis, these results indicate that partial reprogramming alleviates senescence-associated mitochondrial anti-apoptotic states.

At the transcriptional level, OSKM and 2c induced distinct remodeling programs with limited overlap in fully reverted genes. Both interventions suppressed mesenchymal drift, a transcriptional feature implicated in aging and fibrosis^50^. However, OSKM expression induced broader identity remodeling, including partial suppression of fibroblast-specific programs and induction of pluripotency-associated signatures, whereas 2c selectively attenuated mesenchymal drift and TGFβ-associated signatures without detectable loss of somatic identity. This selectivity likely reflects, at least in part, direct inhibition of TGFβ signaling by RepSox^56,57^, and epigenetic modulation by TCP^58^, suggesting that chemical partial reprogramming may achieve senomorphic benefits with a more restricted and through a potentially safer route.

Building on these mechanistic insights, we explored the impact of 2c treatment in very old mice. Administration of 2c to 29-month-old animals improved motor performance, strength, and cognitive parameters, accompanied by reduced expression of senescence- and inflammation-associated genes across multiple tissues and decreased circulating levels of age-associated inflammatory factors, many of which are SASP components. These improvements occurred without detectable loss of SAβGal-positive cells, supporting the conclusion that 2c primarily modulates senescent cell function rather than eliminating senescent cells.

Collectively, our work demonstrates that both genetic and chemical partial reprogramming exert potent senomorphic effects by suppressing SASP abundance and activity, restoring mitochondrial function and apoptotic priming, and improving functional and inflammatory parameters in aged organisms, all while preserving the stable proliferative arrest characteristic of senescence. These findings reveal that key detrimental outputs of senescent cells can be uncoupled from irreversible cell-cycle arrest and identify selective chemical partial reprogramming as a promising safer strategy to mitigate senescence-driven aging phenotypes without erasing cell identity.

## Data availability

All sequencing data generated and analyzed in this study have been deposited in the NCBI Gene Expression Omnibus (GEO) under accession number GSE311930. This includes all raw and processed bulk RNA-sequencing data used in the analyses. Source data underlying all figures and supplementary tables are provided with this manuscript. All other data supporting the findings of this study are available from the corresponding author upon reasonable request.

## Acknowledgements

Work in the laboratory of Manuel Collado is funded by grants from MCINN/AEI/FEDER, UE (PID2021-125479OB-I00), GAIN, Xunta de Galicia (IN607B2024/13) and The Foresight Institute. Victor Nuñez-Quintela was supported by a predoctoral fellowship from GAIN, Xunta de Galicia (IN606A-2021/016). Miguel Angel Prados was supported by predoctoral fellowships from GAIN, Xunta de Galicia (ED481A2022/432) and FPU program from MEC (FPU23/02493). Jeremy Chantrel was supported by DIM Longévité et vieillissement PhD fellowship, LabEx Revive, and FRM 4^th^ PhD fellowship. Han Li’s laboratory is funded by Institut Pasteur, Centre National pour la Recherche Scientific and the Agence Nationale de la Recherche (Laboratoire d’Excellence Revive, Investissement d’Avenir; ANR-10-LABX-73). This work is supported by Agence Nationale de la Recherche (ANR-21-CE13-0006-01). Work in the HL laboratory was also funded by Agence Nationale de la Recherche (ANR-16-CE13-0017, ANR-22-CE16-0015-03, ANR-25-CE13-0056-01), Foundation ARC (PJA PJA 20161205028, 20181208231), and AFMTELETHON (22403).

## Author Contributions

HL and MC conceptualized and supervised the study. VN-Q, JC, MAP performed most of the experiments with the help of PP, IL, LL-D, CC, RP, AF-F, DG-P, AM-D, SDS-A, RL-B, MG-B, CA, JM, PM, DC, FP, MS, MK and AG-D. HL, MC, VN-Q, JC and MAP analyzed the results and prepared the figures. HL and MC wrote the manuscript, and all the authors revised the manuscript and provided feedback.

## METHODS

### ANIMALS AND ETHICAL APPROVALS

All animal procedures were performed in accordance with institutional and national guidelines and approved by the relevant animal welfare committees at each participating institution. Animals were maintained under specific-pathogen-free (SPF) conditions with a 12 h:12 h light:dark cycle and room temperature 20–24 °C unless indicated otherwise.

### i4F-A mice (CEBEGA, Universidade de Santiago de Compostela)

Inducible reprogrammable i4F-A mice (C57BL/6J background) carrying a TetO-driven polycistronic OSKM cassette at the Neto2 locus and rtTA at the Rosa26 locus were used (Abad et al., 2013). The colony was maintained homozygous. Genotypes were confirmed by PCR (Supplementary Table 1). Mice received doxycycline hyclate (doxycycline, Fagron Ibérica) in drinking water at 0.2 mg·mL⁻¹ supplemented with 7.5% sucrose for 2.5 weeks.

### HET3 aged cohort (CEBEGA, Universidade de Santiago de Compostela)

Female HET3 mice, 29 months old, were used for systemic 2c treatment studies (n = 8 control, n = 8 treated). Body weight was recorded weekly and blood glucose measured after a 6-h fast (Accu-chek, Roche).

### Mice used for BH3 profiling (IRB Barcelona)

Animal experiments performed at IRB Barcelona were approved by the institutional ethics committee (Science Park of Barcelona) and conducted in SPF housing with 12 h:12 h light:dark cycles, ambient 20–24 °C and 30–70% humidity. Animals were fed ad libitum with SAFE R40 pellet diet (Safe).

### Euthanasia and tissue handling

Animals were euthanized by CO₂ asphyxiation followed by cervical dislocation. Organs for histology were fixed in 10% neutral-buffered formalin; tissues for molecular analyses were snap-frozen in liquid nitrogen and stored at −80 °C.

### IN VIVO PROCEDURES

#### In vivo reprogramming with conditioned media

Producer cells (5×10⁵) were incubated for 24 h in serum-free medium; supernatants were clarified by 0.45 μm filtration (PVDF filter) and desalted using PD MidiTrap G-10 columns (GE Healthcare, 28-9180-10). Conditioned media (CM) were lyophilized overnight on a LyoQuest 55 (Telstar), reconstituted in 100 μL PBS immediately before use and administered intraperitoneally (IP) on days 1, 3 and 5 (three alternating injections corresponding to CM from a total of 1.5×10⁶ cells per mouse). Lyophilized material was stored short-term at −20 °C if not used immediately.

### Rotarod

Animals underwent three training sessions (5 rpm, 60 s) with 5 min inter-trial intervals. Test trials used a rotarod accelerating from 4 to 40 rpm over 300 s. Latency to fall (or passive rotation completion) was recorded. Three trials per animal were performed with 10 min rest between trials. (Ugo Basile Rotarod)

### Grip hanging test

Mice were placed on a wire mesh which was gently inverted to start the trial. Each mouse performed three trials with ∼1 min rest between trials.

### Open field

Animals were placed in an open-field arena and allowed to explore for 60 min. Sessions were recorded from above and analyzed using AnyMaze tracking software (Stoelting Co.). Metrics included total distance travelled, maximum speed and number of center entries.

### Y-maze

Spontaneous alternations were assessed over an 8 min free-exploration period in a Y-shaped maze. Sessions were recorded and analyzed with AnyMaze.

### Elevated zero maze

The circular elevated zero maze comprised two open arms and two closed arms. Each mouse started in a closed arm and was allowed to explore; time in open arms, distance travelled and open-arm crossings were quantified.

## CELL CULTURE, PRIMARY CELL DERIVATION AND LINES

All cells were cultured at 37 °C in a humidified 5% CO₂ incubator and handled under aseptic conditions. Mycoplasma testing by PCR was performed routinely.

### Media

Standard medium: high-glucose DMEM (4,500 mg·L⁻¹; Corning, 10-013-CV) supplemented with 10% fetal bovine serum (FBS; Corning, 35-010-CV), 2 mM L-glutamine and 1% penicillin/streptomycin (Gibco, 15140-122). Reprogramming medium: high-glucose DMEM supplemented with 15% KnockOut Serum Replacement (KSR; Gibco, 10828028), 2 mM L-glutamine, 1% penicillin/streptomycin, 1% non-essential amino acids (Gibco, 11140-050), 0.2% β-mercaptoethanol (Gibco, 21985-023) and 0.2% LIF (Gibco, ESG1107).

### Primary MEFs

Mouse embryonic fibroblasts (MEFs) were derived independently from E13.5 embryos (C57BL/6J WT and i4F-A). After removal of head and viscera, embryonic tissues were minced and digested in 1.5 mL trypsin-EDTA (0.25%, Gibco, 25200-072) at 37 °C for 10 min, mechanically dissociated, resuspended in standard medium and expanded. Cells were cryopreserved in freezing medium (standard medium + 10% DMSO) at 4×10⁶ cells per vial, held at −80 °C 24 h and transferred to liquid nitrogen for long-term storage. Unless otherwise indicated, all experiments using MEFs were performed with a minimum of three independent biological replicates (distinct embryo-derived MEF lines), with at least two technical replicates per experiment.

### Cell lines

HEK293T and RAW 264.7 cells were obtained from ATCC and cultured in standard medium. Human dermal fibroblasts immortalized with hTERT carrying a doxycycline-inducible GFP-Progerin construct (kindly donated by Dr Tom Misteli^29^) were cultured in standard medium.

### Conditioned media production (in vitro) and protein profiling

Producer cells were plated and 12 h later medium was replaced; cells were incubated 48 h to condition the medium. CM were centrifuged 500 g for 10 min, filtered (0.45 μm) and used fresh or stored at 4 °C for short periods. Secreted proteins were profiled using the Proteome Profiler Mouse XL Cytokine Array (R&D Systems, ARY028; confirm catalog number in final version). Membranes were developed and imaged on a ChemiDoc MP Imaging System (Bio-Rad, Model: ChemiDoc MP, Cat. #1708280) and quantified using the Protein Array Analyzer macro in ImageJ/Fiji (NIH).

### Viral production and transduction

Lentiviral particles were produced in HEK293T cells. Packaging plasmids (ViraPower™ Lentiviral Packaging Mix, Invitrogen, K497500) and plasmid of interest were co-transfected using PEI (Polysciences, 23966-2), with the ratio of DNA:PEI 1:6. Viral supernatants were collected 24 h after transfection, filtered through 0.45 μm PVDF filters, supplemented with hexadimethrine bromide (Polybrene; Sigma-Aldrich, TR-1003-G) at 8 μg·mL⁻¹ and applied to target cells for 12 h. Infection was repeated for three rounds. Plasmids used: Tet-O-FUW-OSKM^59^ (Addgene #20321), FUW-M2rtTA^60^ (Addgene #20342), TetO-FUW-Oct4^61^ (Addgene #20323), TetO-FUW-Sox2^61^ (Addgene #20326), TetO-FUW-Klf4^61^ (Addgene #20322), and pBABE puro H-Ras V12 (Addgene #9051, a gift from William Hahn). Plasmid amplification used Stbl3 competent E. coli (Thermo Fisher Scientific) and midipreps (NucleoBond Xtra Midi, Macherey-Nagel, 740410.50). DNA concentration was measured on a NanoDrop One spectrophotometer (Thermo Fisher Scientific, ND-ONE-W).

### Doxycycline treatment (in vitro)

Doxycycline (Sigma-Aldrich, D9891) was used at 5 μg·mL⁻¹ to activate rtTA/TetO systems in cell culture; medium and doxycycline were refreshed every 48 h.

### 2c in vitro

RepSox and trans-2-phenylcyclopropylamine hydrochloride were used at 10 μM each for 6 days; media and compounds were refreshed every 48 h.

### Irradiation-induced senescence

Senescence was induced by X-ray irradiation using a CLINAC 2300 IX linear accelerator (Varian Medical Systems, CLINAC 2300 IX) at a total dose of 10 Gy delivered at 5.2 Gy·min⁻¹ in two opposing fields (0° and 180°). Medium was refreshed 2 h post-irradiation.

### Adipose tissue dissociation and adipocyte isolation

Visceral white adipose tissue was dissected into cold HBSS (Gibco, 14025-092), minced and incubated 1 h at 37 °C with agitation in digestion buffer containing DNase I (125 U; Sigma-Aldrich, DN25), hyaluronidase (100 U; Sigma-Aldrich, H3506), collagenase IV (300 U; Thermo Fisher, 17104-019) and 2% FBS. Samples were filtered through a 100 μm mesh and centrifuged at 150 g for 8 min. The top adipocyte layer was collected, washed in HBSS and centrifuged again at 150 g for 8 min prior to BH3 profiling.

### BH3 assay

BH3 profiling was performed as previously described^44,62^ with adaptations for adipocytes and MEFs. Cells were stained with Zombie Violet viability dye (BioLegend, 423113) and resuspended in MEB buffer (150 mM mannitol, 10 mM HEPES–KOH pH 7.5, 150 mM KCl, 1 mM EGTA, 1 mM EDTA, 0.1% BSA, 5 mM succinate). A 25 μL cell suspension was mixed with 25 μL BIM BH3 peptide (serial final concentrations: 0.01, 0.03, 0.1, 0.3, 1, 3, 10, 30, 100 μM) prepared in MEB with 0.002% digitonin and incubated in 96-well plates (Corning, 3795) for 1 h at room temperature. Samples were fixed in 4% formaldehyde, permeabilized and stained for cytochrome c (Alexa Fluor 647 anti-Cytochrome c, BioLegend, 612310) and p21 (mouse anti-p21, Santa Cruz Biotechnology, sc-6246). Flow cytometric acquisition was performed on a CytoFLEX (Beckman Coulter, Model CytoFLEX S) and analyses conducted in FlowJo (version to be specified). Each condition was assayed at least in triplicate and Δ% priming was calculated as (% cytochrome c released in treated wells) − (% cytochrome c released in untreated wells).

### Population doublings

Population doublings were calculated as n = (log Xf − log Xi)/log 2. Cell counts were obtained using a LUNA-II automated cell counter (Logos Biosystems, LUNA-II).

### EdU/Ki67

For proliferation analysis, cells were incubated with 10 μM EdU (Click-iT Plus EdU Alexa Fluor-647 kit, Invitrogen, C10640) for 4 h, fixed in 3.7% paraformaldehyde, permeabilized with 0.5% Triton X-100 and processed according to the manufacturer’s instructions. Ki67 immunostaining used rabbit anti-Ki67 (SP6, Invitrogen, MA5-14520) at 1:250 in 3% BSA for 1 h at room temperature with Alexa Fluor-488 secondary (Invitrogen, A-11034) at 1:1000. Nuclei were counterstained with Hoechst 33342 (Thermo Fisher Scientific, H3570) at 1 μg·mL⁻¹. Confocal images were acquired on a Leica SP8 confocal microscope (Leica Microsystems, Model SP8).

### Transwell migration

RAW 264.7 migration assays used 8 μm pore transwell inserts (Corning, 3422) in 24-well plates. Lower wells contained 5×10⁴ producer cells in standard medium (10% FBS); 2×10⁴ RAW 264.7 cells in medium with 0.05% FBS were added to the insert. After 48 h, migrated cells were fixed with 3.7% PFA (2 min), permeabilized with 100% methanol (20 min), stained with 0.05% crystal violet (10 min), washed and imaged by brightfield microscopy. Crystal violet was eluted with 33% acetic acid for 5 min with shaking and absorbance measured at 570 nm on a BioTek Epoch2 microplate reader (Agilent/BioTek, Epoch2).

### Mitochondrial dyes and flow cytometry

MitoTracker Green FM (Invitrogen, M7514) was used at 50 nM in HBSS for 30 min to estimate mitochondrial mass. Mitochondrial membrane potential was measured using TMRM (Invitrogen, T668) at 100 nM for 30 min. Mitochondrial ROS were measured with MitoSOX Red (Invitrogen, M36008) at 5 μM for 30 min. After staining, cells were washed, trypsinized, pelleted (1,400 g, 5 min), resuspended in HBSS + 0.01% BSA and analyzed on a CytoFLEX cytometer (Beckman Coulter). At least 5×10⁴ events were recorded per sample. Emission channels used: 516 nm (MitoTracker), 574 nm (TMRM) and 610 nm (MitoSOX).

### Seahorse metabolic flux

OCR and ECAR were measured on a Seahorse XFe96 Analyzer (Agilent Technologies, Model XFe96). Cells were equilibrated in Seahorse XF base medium (pH 7.4) supplemented with glucose (4.5 g·L⁻¹), 1 mM sodium pyruvate and 3.9 mM L-glutamine. Mitochondrial stress test reagents: oligomycin (1 μM), FCCP (1.5 μM) and rotenone + antimycin A (1 μM each). Data were normalized to total protein content per well (BCA assay; Thermo Fisher Scientific, Pierce BCA Protein Assay Kit, 23225) and analyzed with Wave software (Agilent).

### ATP assay

Mitochondrial ATP was determined using CellTiter-Glo (Promega, G7572). Cells were seeded at 5×10⁴ per well in white opaque 96-well plates and incubated 14 h. After a PBS wash, cells were incubated 2 h in metabolic buffer (156 mM NaCl, 3 mM KCl, 2 mM MgSO₄, 1.25 mM KH₂PO₄, 2 mM CaCl₂, 20 mM HEPES) containing 5 mM 2-deoxy-D-glucose and 1 mM pyruvate. Ninety microlitres were removed and mixed with 100 μL CellTiter-Glo, plates were shaken 2 min and luminescence read after 10 min stabilization on an appropriate plate reader (BioTek Epoch2).

### Protein extraction

Cells were lysed in RIPA buffer (150 mM NaCl, 10 mM Tris-HCl pH 7.5, 0.1% SDS, 1% Triton X-100, 5 mM EDTA pH 8.0, 1% sodium deoxycholate) supplemented with protease and phosphatase inhibitors (1 mM Na₃VO₄, 1 mM PMSF, 1 mM DTT, 4 mM NaF and commercial protease inhibitor cocktail; Sigma-Aldrich, P8340). Lysates were clarified by centrifugation and protein quantified by Bradford assay.

### SDS-PAGE and transfer

30 µg protein per lane were denatured, resolved on Bolt 4–12% Bis-Tris gels (Invitrogen, NW04120BOX) and transferred to 0.45 μm PVDF membranes (Merck Millipore, IPVH00010).

### Blotting and detection

Membranes were blocked in 5% BSA in TTBS for 1 h, incubated with primary antibodies overnight at 4 °C, washed and incubated with HRP-conjugated secondary antibodies for 1 h. Detection used SuperSignal West Pico PLUS Chemiluminescent Substrate (Thermo Fisher Scientific, 34580) and images captured on a ChemiDoc MP (Bio-Rad). Information relating to the primary antibodies used is in Supplementary Table 2. HRP anti-mouse and HRP anti-rabbit secondaries (Santa Cruz) were used at 1:10,000.

### Cytokine array

Secreted protein profiling from conditioned media was performed with the Proteome Profiler Mouse XL Cytokine Array (R&D Systems). Membranes were processed according to manufacturer instructions, imaged on a ChemiDoc MP (Bio-Rad) and quantified using ImageJ with the Protein Array Analyzer macro.

### Immunofluorescence and microscopy

Cells were fixed in 3.7% paraformaldehyde, permeabilized with 0.5% Triton X-100 and blocked in 3% BSA. Primary antibodies were incubated 1 h at room temperature followed by fluorescent secondary antibodies (Alexa Fluor conjugates, Invitrogen). γH2AX (Ser139; Cell Signaling Technology, 9718, 1:500) staining was performed as an example of DNA damage staining. Nuclei were counterstained with DAPI (Thermo Fisher Scientific, D1306) at 0.1 μg·mL⁻¹. Confocal images were acquired on a Leica SP8 confocal laser scanning microscope (Leica Microsystems, SP8) using LAS X.

### SAβGal histochemistry

Cells were fixed in 2% paraformaldehyde + 0.2% glutaraldehyde for 15 min, washed and incubated at 37 °C (CO₂-free) for 8 h in staining solution containing 40 mM citric acid/sodium phosphate buffer pH 5.6, 150 mM NaCl, 5 mM K₃Fe(CN)₆, 5 mM K₄Fe(CN)₆, 2 mM MgCl₂ and 1 mg·mL⁻¹ X-Gal (5-bromo-4-chloro-3-indolyl β-D-galactopyranoside). Images were captured at 10× and positive cells quantified using ImageJ.

### Flow cytometric SAβGal detection

CellEvent™ Senescence Green Assay Kit (Invitrogen; C10840) was used according to the manufacturer’s instructions. Samples were analyzed on a CytoFLEX and 1×10⁴ events per sample recorded.

## RNA EXTRACTION, QPCR AND RNA-SEQ

### RNA extraction

Total RNA from cultured cells was isolated with NucleoSpin RNA Plus kits (Macherey-Nagel, 740984.50) following the manufacturer’s instructions. Tissue RNA was extracted in TRIzol Reagent (Thermo Fisher Scientific, 15596026) with mechanical homogenization (TissueLyser II, Qiagen, 85300) followed by chloroform extraction and isopropanol precipitation; pellets were washed with 75% ethanol and resuspended in RNase-free water. RNA quality and integrity were assessed on an Agilent 4200 TapeStation (Agilent Technologies); samples with RIN ≥ 8 and 28S/18S ≥ 1 were used for downstream library preparation.

### RT–qPCR

cDNA was synthesized using the High-Capacity cDNA Reverse Transcription Kit (Applied Biosystems, 4368813) on a MultiGene OptiMax thermocycler (Labnet). qPCR reactions were run on a QuantStudio 3 Real-Time PCR System (Thermo Fisher Scientific, A28137) using NZYSpeedy qPCR Green Master Mix ROX (NZYTech, MB12501). Each 20 μL reaction contained 33 ng cDNA and 0.25 μM forward and reverse primers; reactions were performed in triplicate and normalized to *Wdr55*. qPCR data were analyzed with QuantStudio Design & Analysis Software v2.8. See primers sequences at Supplementary Table 3.

## RNA-SEQ AND BIOINFORMATIC ANALYSIS

### Raw data quality control and preprocessing

Libraries were prepared and sequenced by BGI (DNBSeq, paired-end 100 bp). Raw.fastq files were subjected to standard quality assessment and filtering to remove low-quality bases and technical artefacts using SOAPnuke^63^. This step was conducted by the Beijing Genomics Institute (BGI) company using the following parameters: -n 0.001 -l 20 -q 0.4 --adaMR 0.25 --polyX 50 --minReadLen 100. Reports are present in Supplementary Table 4.

### Read alignment and quantification

Cleaned reads were aligned to the most up-to-date mouse reference genome (GRCm39) using the STAR^64^ (2.7.10b) splice-aware aligner with the following parameters: --runMode alignReads --outSAMtype BAM SortedByCoordinate. Reports are present in Supplementary Table 4. Then, gene-level read count matrices were generated using featureCounts^65^ from the Subread^66^ package, relying on an up-to-date gene annotation corresponding to the reference genome (GRCm39).

### Differential expression analysis

All subsequent analyses were performed in R (v4.4.1). For each dataset (OSKM and 2c), read count matrices were imported and processed independently using the DESeq2^67^ package. Lowly expressed genes were filtered out according to DESeq2’s standard recommendations (*i.e.*, genes with reads < 10 counts for a minimal number of 3 samples were screened out). Size-factor estimation, dispersion computation, and negative binomial modelling were performed following the canonical DESeq2 pipeline. Differential expression contrasts were computed individually using the default Wald test for each biological comparison (*e.g.*, senescent *vs.* proliferating cells; OSKM-expressing senescent *vs.* senescent cells; 2c-treated senescent *vs.* senescent cells, *etc.*). Significant Differentially Expressed Genes (DEGs) were determined by an *a priori* alpha risk of 5% and an *a priori* threshold of |log2FC| > 1 (Supplementary Table 5). Resulting p-values were then adjusted using the Benjamini-Hochberg’s method^68^ for False Discovery Rate (FDR). Histograms of p-values for each comparison were verified to follow the expected pattern: *i.e.*, uniform distribution except an expected peak of significance < 0.05 and another one for significance > 0.90 (Supplementary Table 4). For data visualization (*e.g.*, MA^69^ and volcano plots), shrinkage of the log_2_ fold changes (log_2_FC) was performed using the ashr^70^ method according to DESeq2 guidelines.

### Data transformation and exploratory analyses

Variance-stabilising transformation (vst) was applied to normalised count data, and were used as inputs for visualisation (*e.g.*, heatmaps), clustering (*i.e.*, PCA), pattern plots (*i.e.*, via the DEGreport package^71^), and pathway enrichment analysis (*i.e.*, via GSEA). Principal component analysis (PCA) and sample-to-sample distance heatmaps were generated to assess batch structure, replicate clustering, and overall transcriptional variation across conditions. PCA was computed using the plotPCA() function from the DESeq2 package. In brief, this function uses the standard prcomp() function on the top 500 most variable genes in the 3 biological conditions of interest (3 replicates per condition) to calculate the usual PCA metrics. The resulting first two principal components (PC) were used to visualize sample clustering and to identify major sources of variance across experimental conditions. Volcano plots, heatmaps, and other graphical summaries were generated using ggplot2^69^ and associated packages. For heatmaps, values are scaled per gene (row-wise Z-scores) and indicate relative upregulation (white-to-red) and downregulation (white-to-blue), respectively. Rows were ordered by unsupervised clustering to regroup genes with similar inter- and intra-condition variability.

### Pathway enrichment analyses

To investigate functional enrichment in particular pathways in each of the comparisons of interest, Gene Set Enrichment Analyses (GSEA) were performed using the vst-transformed normalized count data using the official GSEA software^72,73^ (v4.3.2) on a subset of gene sets prior to the test (Supplementary Table 4). 10 000 permutations using the “gene_set” permutation type parameter were performed for these two groups of gene sets separately. Genes sets with an adjusted p-value (q-value_FDR_) < 0.05 were considered significantly different. Enrichment plots were generated by the GSEA software using the parameters described above. To investigate functional enrichment in particular pathways in significant DEGs from each of the comparisons of interest, Over-Representation Analyses (ORA) were performed using the online gprofiler:GOSt^74^ tool on the ranked lists of significant DEGs with Gene Ontology – Biological Process (GOBP) as reference database. DEGs were ranked based on the following formula: log_2_FC x -log_10_(p-value) in order to take into account both the magnitude and the significance of the expression change. The networks of the significant gene sets (q-value_FDR_ < 0.05) were then visualised using the Cytoscape software^75^ with the following default parameters: Jaccard (50%) + Overlap (50%) combined. Clusters were manually annotated and regrouped based on the similarities of the implicated biological processes.

### Reproducibility

All analyses were executed using reproducible R scripts and version-controlled environments to ensure transparency and traceability.

## STATISTICS

Unless otherwise stated, data are presented as mean ± standard deviation (SD). Two-group comparisons used two-tailed Student’s t-tests; comparisons across multiple groups used one-way ANOVA with Tukey’s post-hoc test. Normality was tested using the Shapiro–Wilk test. Outliers were detected with ROUT (Q = 1%) and excluded from downstream analyses where indicated; excluded data points are reported in figure legends. Exact sample sizes (n), definition of biological versus technical replicates and exact statistical tests are specified in figure legends. For experiments using MEFs, biological replicates correspond to independent MEF lines derived from distinct embryos, and technical replicates refer to repeated measurements within the same biological sample.

---

**Extended Data Fig 1.**
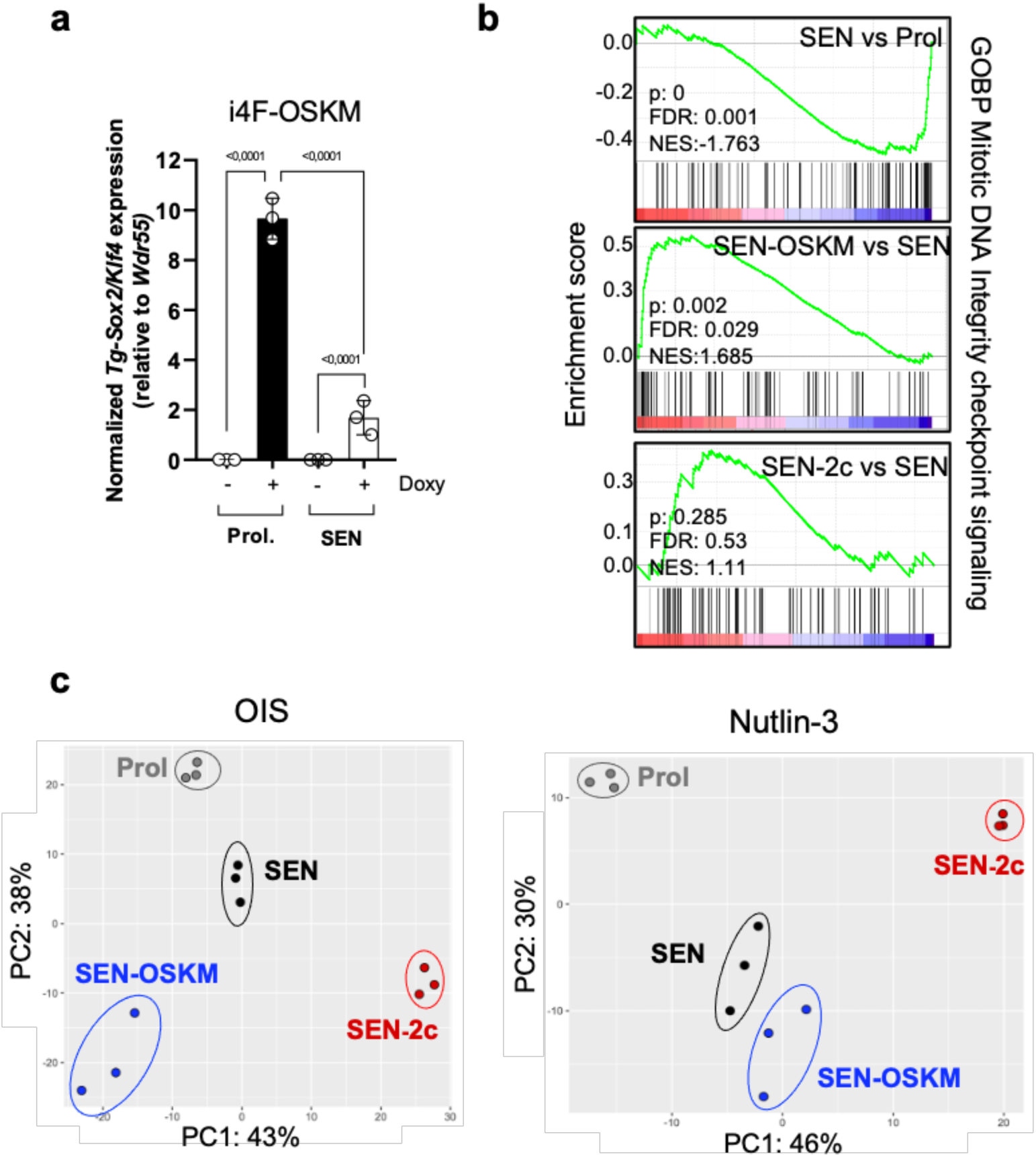
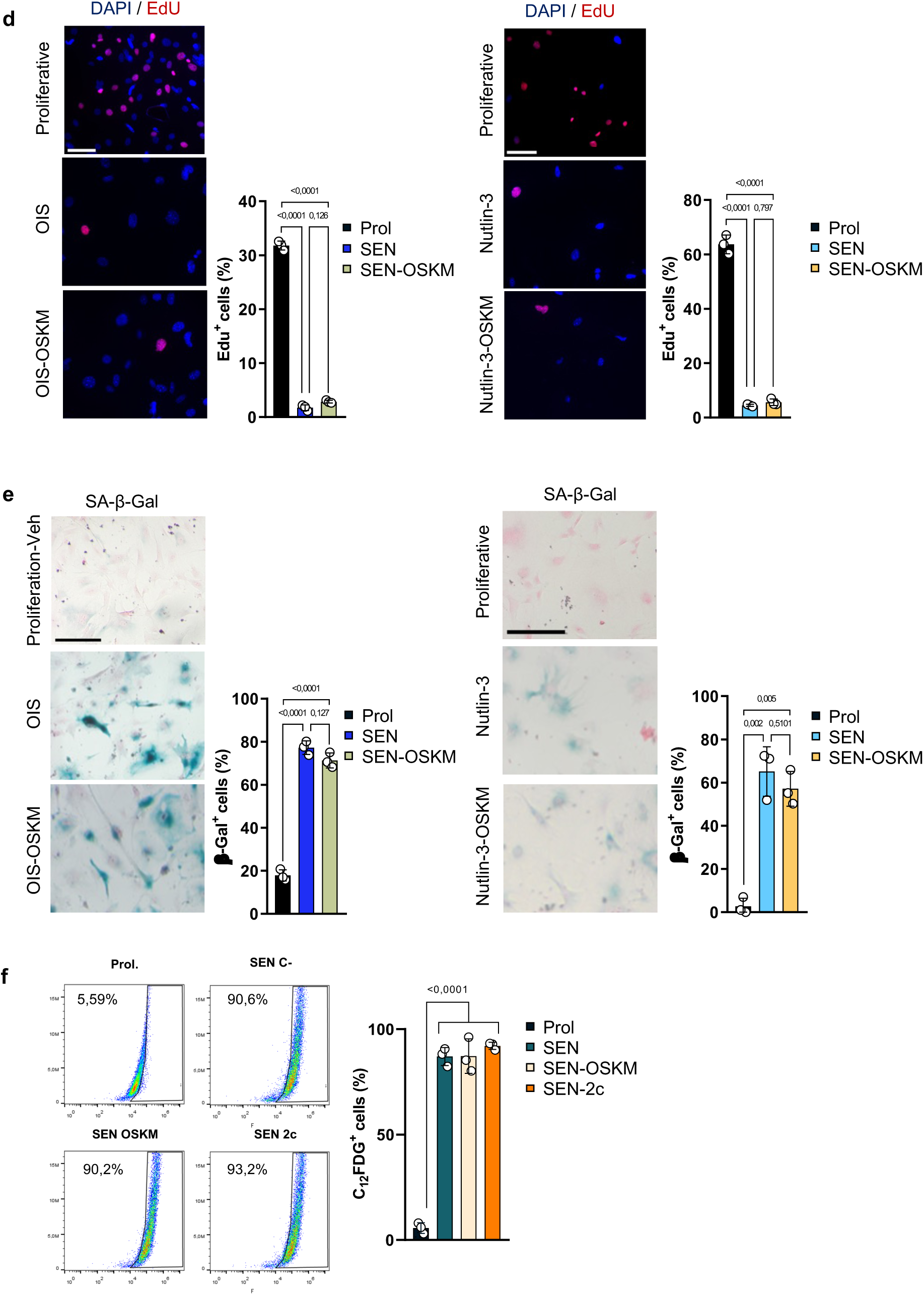
Effects of OSKM expression or 2c treatment across distinct senescence models. **a,** RT–qPCR analysis of OSKM cassette expression (2^ΔΔCt^) in transgenic i4F MEFs in the presence or absence of doxycycline (Doxy). **b,** Gene set enrichment analysis of the “Mitotic DNA Integrity checkpoint signaling” gene set. **c,** Principal component analysis of bulk RNA-seq data from oncogene-induced senescent (OIS, left) and nutlin-3–induced senescent MEFs (right). **d,** Representative immunofluorescence images of EdU incorporation and corresponding quantification in OIS (left), and nutlin-3–induced senescent MEFs (right). Nuclei were counterstained with DAPI. **e,** Senescence-associated β-galactosidase activity assessed by X-gal staining in OIS (left) and nutlin-3–induced senescent MEFs (right), with quantification. **f,** Flow cytometry analysis of SAβGal activity on irradiated MEFs, using the fluorescent substrate C_12_FDG, with quantification. (**a**, **d**, **e** and **f**), Statistics were performed using one-way ANOVA followed by Tukey’s post hoc test. Error bars represent the s.d.

**Extended Data Fig 2.**
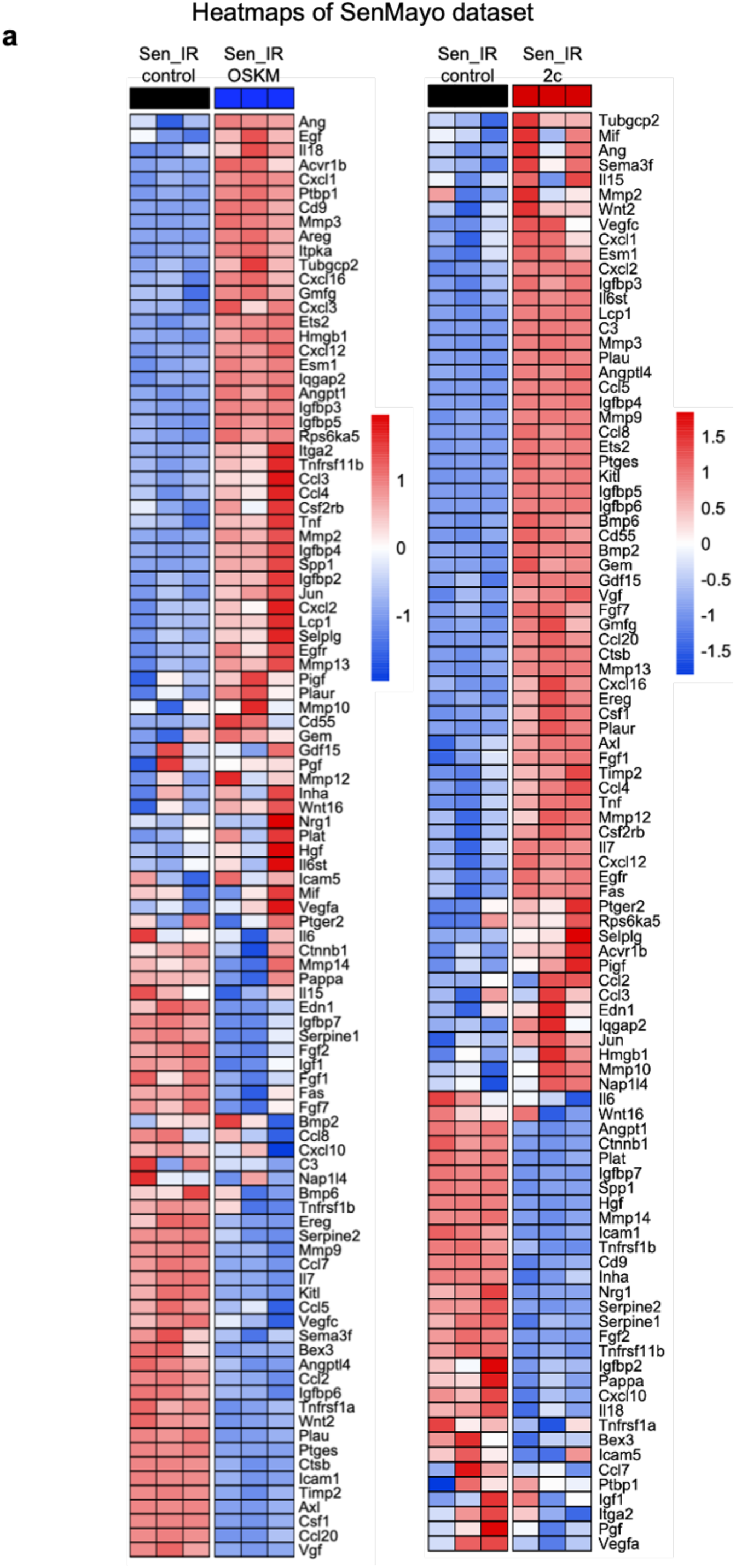
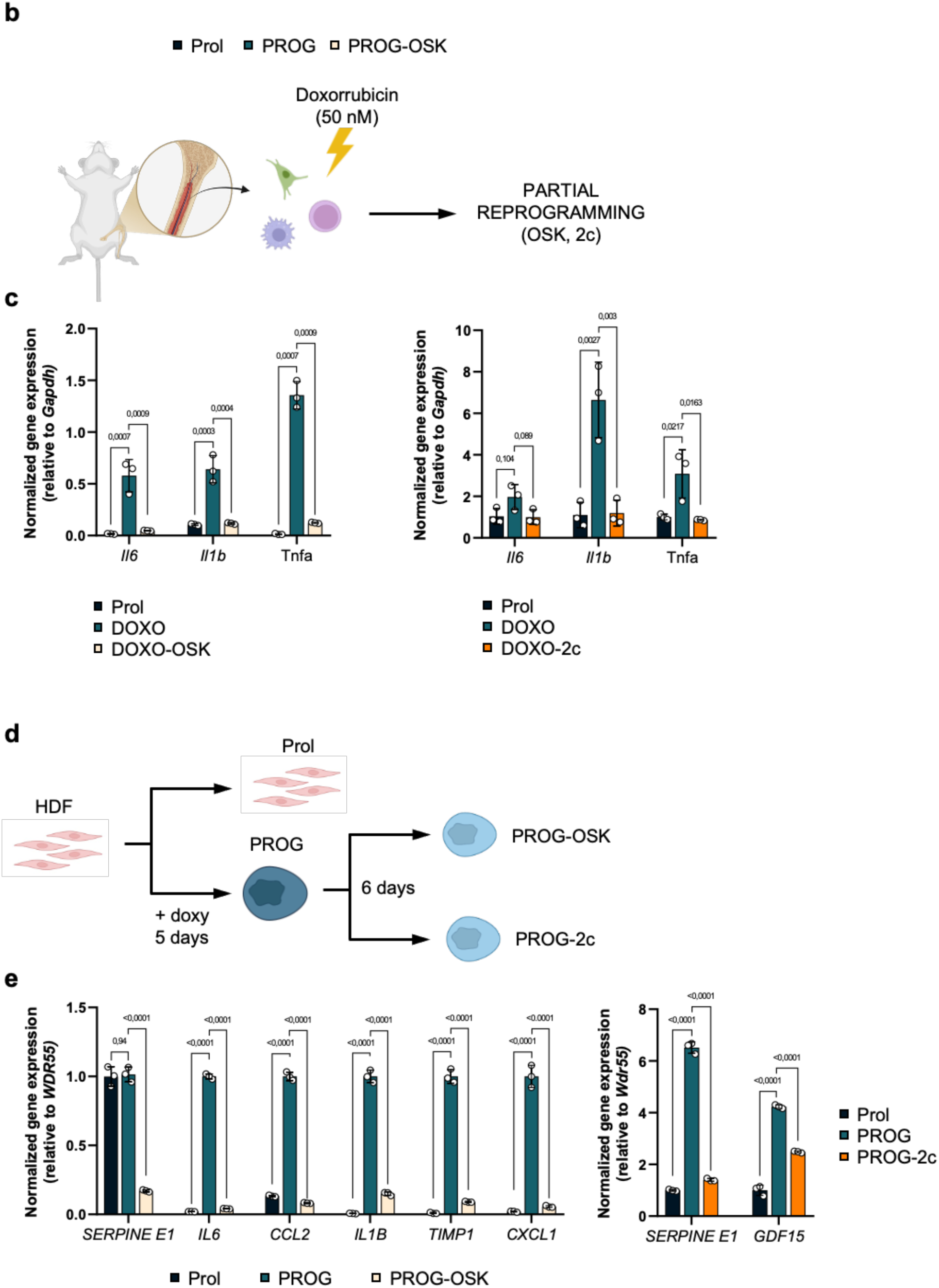
Partial reprogramming modulates senescence phenotypes. **a,** Heatmaps depicting the relative expression profiles of genes belonging to the SenMayo gene signature based on bulk RNA-seq data (Z-score normalized). **b,** Schematic representation of the *ex vivo* experimental design using mouse bone marrow (BM) cells treated with Doxorubicin (Doxo). **c,** RT–qPCR analysis of *Il6*, *Il1b*, and *Tnfa* mRNA levels (2^ΔΔCt^) in BM cells, normalized to *Gapdh*. **d,** Schematic representation of the experimental design using human dermal fibroblasts (HDFs) with inducible progerin (PROG) expression. **e,** RT–qPCR analysis of indicated SASP factor mRNA levels (2^ΔΔCt^) in HDFs, normalized to *Wdr55*. (**c, e**) Statistics were performed using one-way ANOVA followed by Tukey’s post hoc test. Error bars represent the s.d.

**Extended Data Fig 3.**
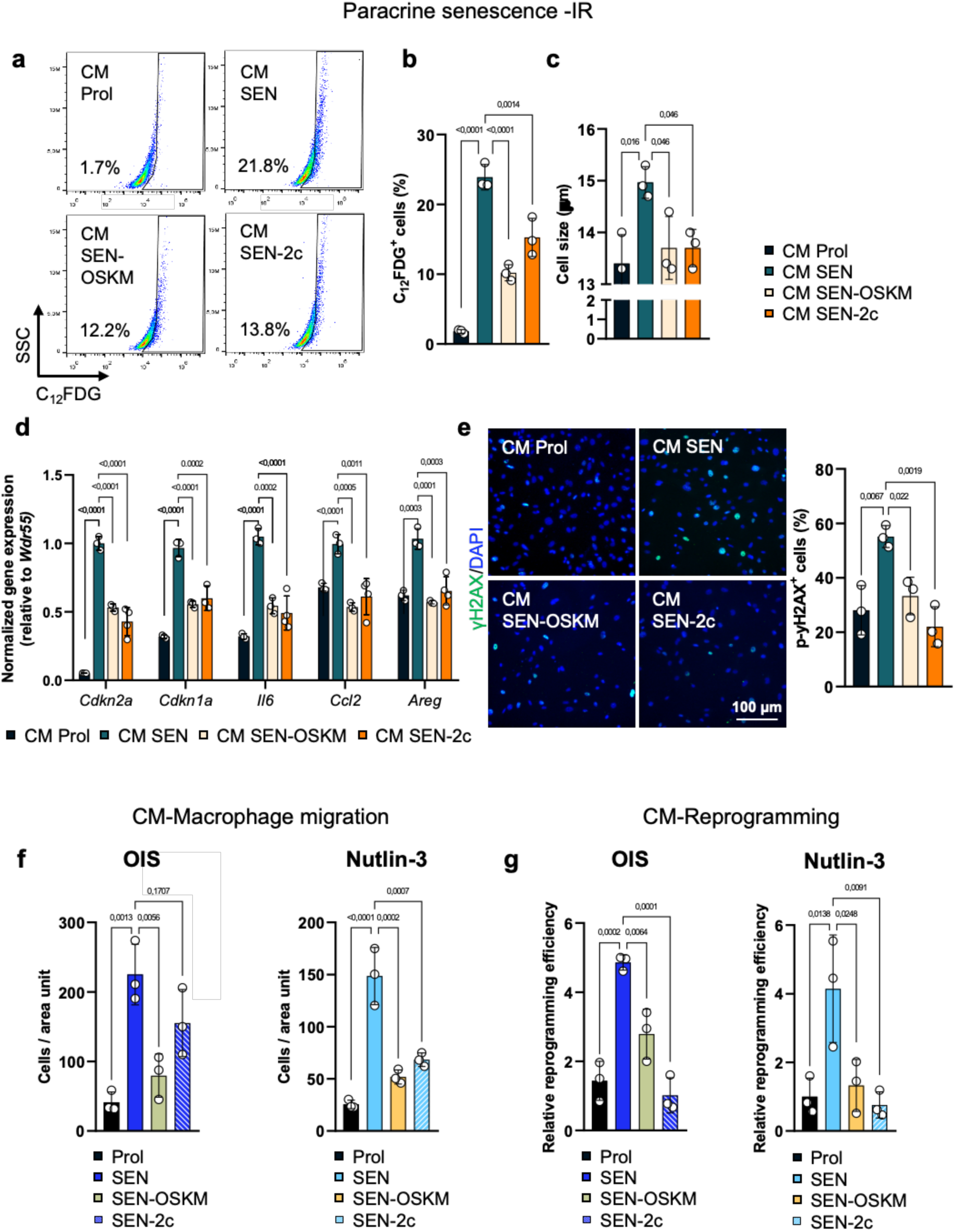
Partial reprogramming attenuates paracrine senescence induction and SASP-mediated activities across different senescence models. **a,b,** Flow cytometry analysis of SAβGal activity in recipient MEFs using C_12_FDG, showing representative plots (**a**) and corresponding quantification (**b**). **c,** Quantification of cell size in recipient MEFs. **d,** RT–qPCR analysis of *Cdkn2a*, *Cdkn1a*, *Il6*, *Ccl2*, and *Areg* mRNA levels (2^ΔΔCt^) in recipient cells normalized to *Wdr55*. **e,** Representative immunofluorescence images (left) and quantification (right) of γH2AX-positive recipient cells. Nuclei were counterstained with DAPI. Scale bar, 100 µm. **f,** Quantification of bone marrow derived macrophage (BMDM) migration toward CM derived from OIS- or nutlin-3-induced senescent cells. **g,** Relative reprogramming efficiency of i4F MEFs cultured with CM from OIS- or Nutlin-3-induced senescent cells. (**b–g**) Statistics were performed using one-way ANOVA followed by Tukey’s post hoc test. Error bars represent the s.d.

**Extended Data Fig 4.**
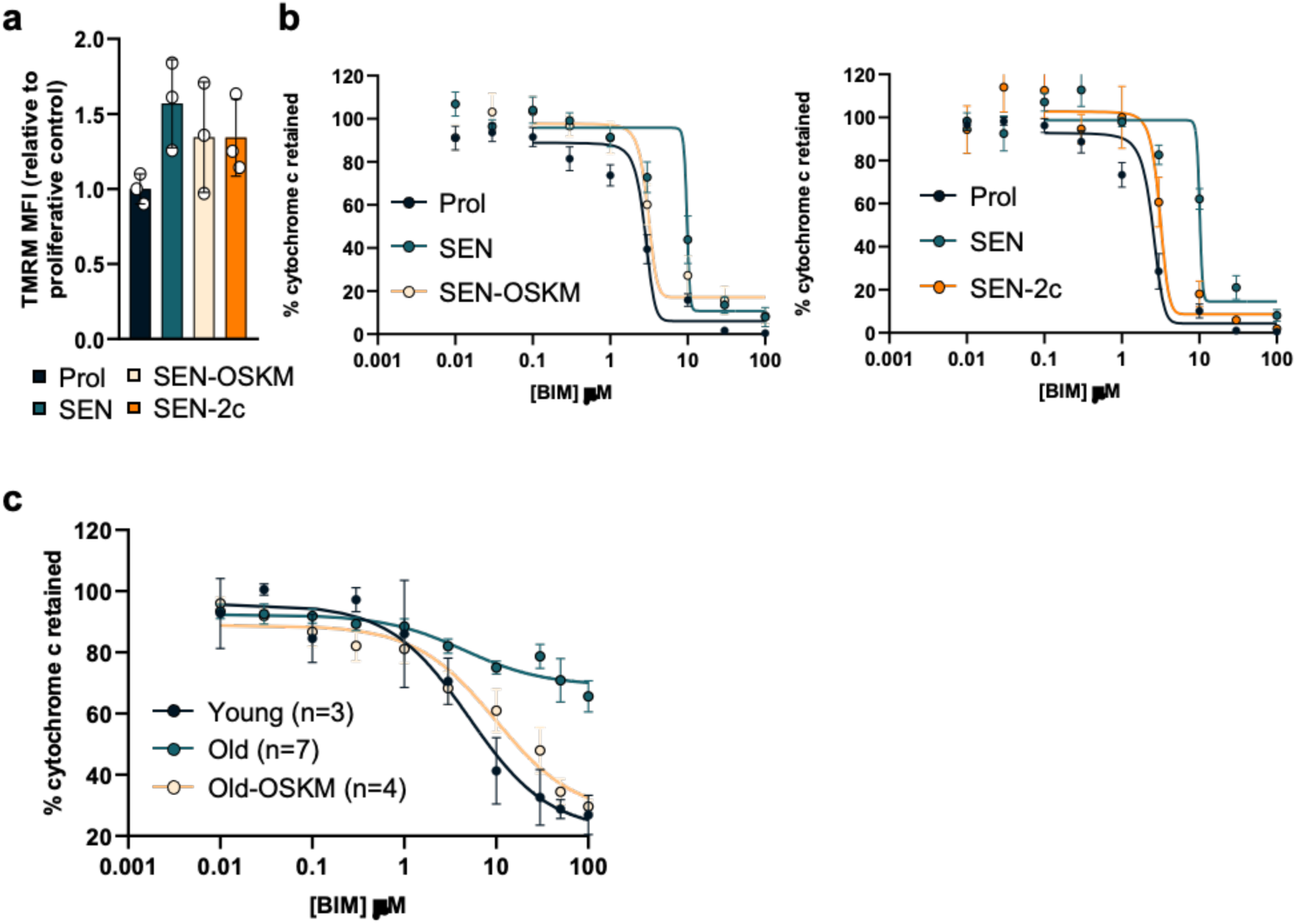
Characterization of mitochondrial membrane potential and apoptotic priming in MEFs and adipose tissue. **a,** Mitochondrial membrane potential assessed by TMRM Mean Fluorescence Intensity (MFI) in MEFs. **b,** Cytochrome c retention curves in response to increasing concentrations of BIM peptide in MEFs following OSKM expression (left) or 2c treatment (right). **c,** Cytochrome c retention curves in response to BIM peptide in adipocytes isolated from visceral white adipose tissue (*n*=3 middle aged, *n*=7 old, *n*=4 old-OSKM). (**a**) Statistics were performed using one-way ANOVA followed by Tukey’s post hoc test. Error bars represent the s.d.

**Extended Data Fig 5.**
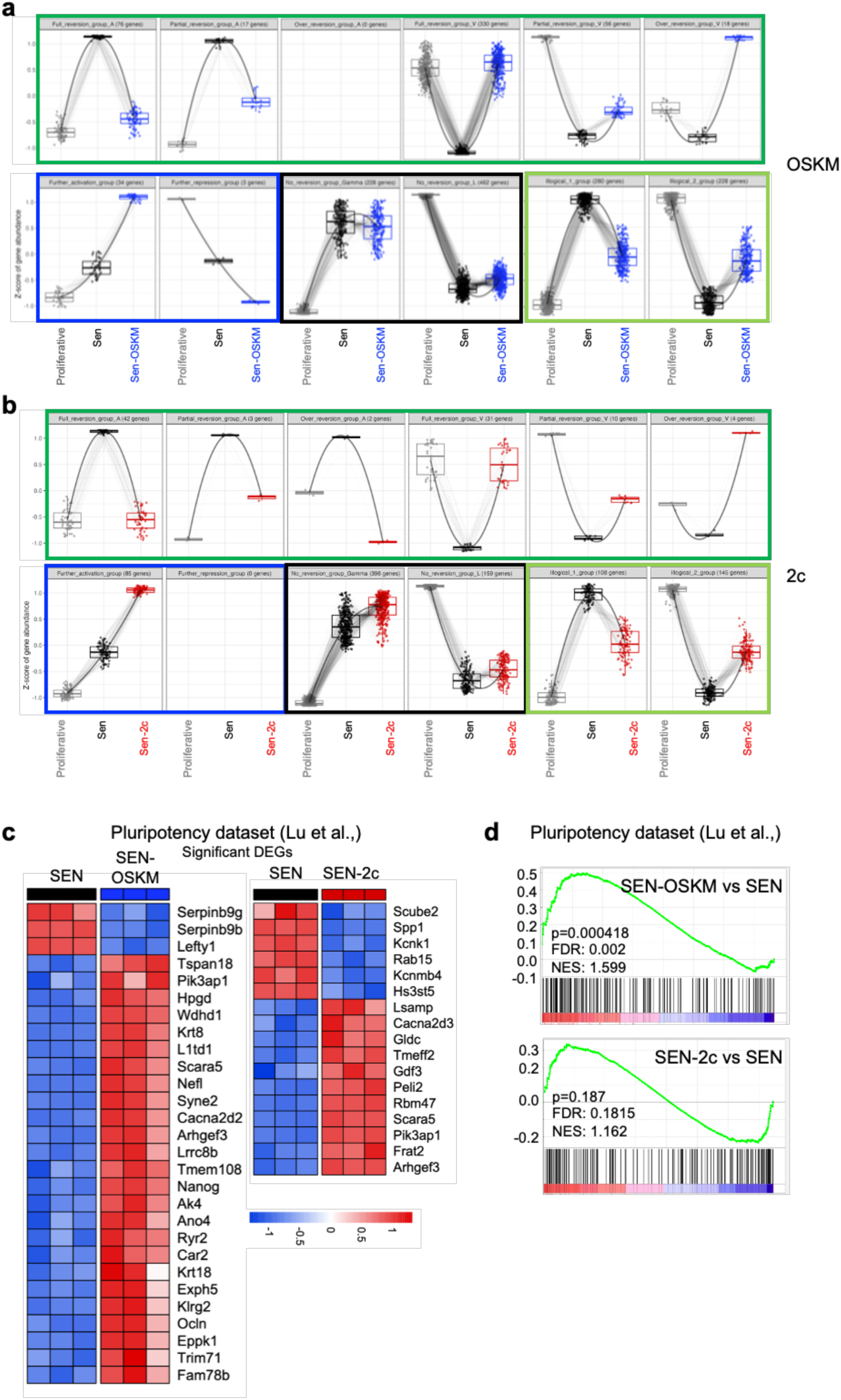
Transcriptional response categories and pluripotency signature analysis. **a,b,** Pattern plots of senescence-associated DEGs categorized by transcriptional response to OSKM expression (**a**) or 2c treatment (**b**). Individual lines represent single genes, and box plots show the distribution of Z-scores across experimental conditions. **c,** Heatmaps of relative expression for significant DEGs belonging to the pluripotency gene signature^50^ in senescent cells compared to OSKM-treated (left) or 2c-treated (right) cells. The color scale indicates Z-score normalized expression levels. **d,** Gene set enrichment analysis of the pluripotency gene signature comparing SEN-OSKM vs. SEN (top) and SEN-2c vs. SEN (bottom).

**Extended Data Fig 6.**
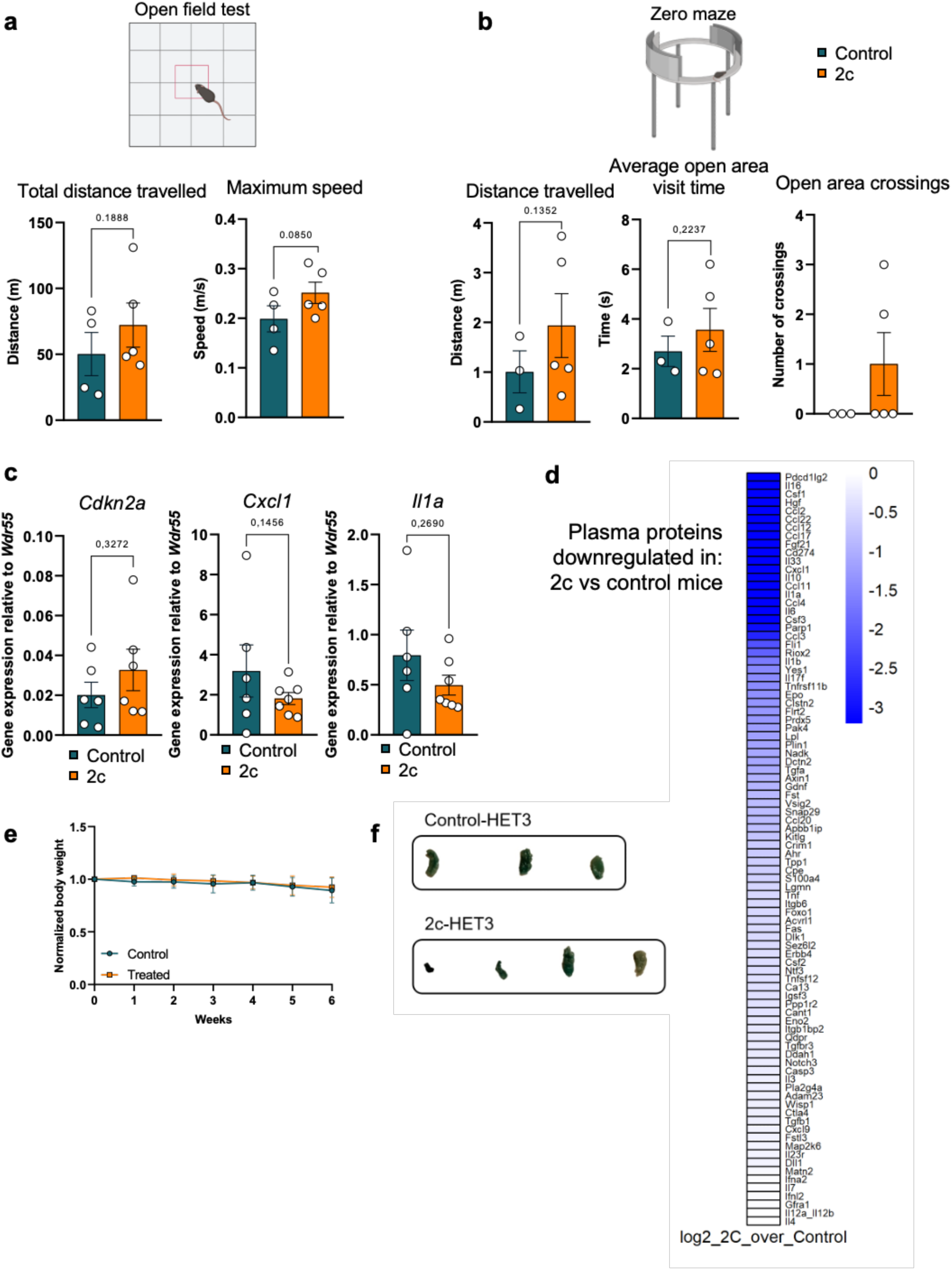
Behavioral and tissue-level characterization of 2c-treated aged mice. **a,** open-field test assessing locomotor activity and anxiety-like behavior, showing total distance travelled (left) and maximum speed (right). **b,** Elevated zero maze performance, including total distance travelled (left), average visit time in the open area (middle), and the number of open area crossings (right). **c,** RT–qPCR analysis of hepatic *Cdkn2a*, *Cxcl1*, and *Il1a* mRNA levels (2^ΔΔCt^) normalized to *Wdr55*. **d,** Heatmap showing the relative abundance (log_2_ fold change) of circulating plasma proteins found downregulated in 2c-treated mice. **e,** Normalized body weight curves over the 6-week treatment period. **f,** SAβGal staining in white adipose tissue from control and 2c-treated HET3 mice. (**a–c**) Statistics were performed using unpaired two-tailed t-test. Error bars represent the s.e.m. Sample size: Control group, *n*=4; 2c group, *n*=5.

## REFERENCES

1 Takahashi, K. & Yamanaka, S. Induction of pluripotent stem cells from mouse embryonic and adult fibroblast cultures by defined factors. Cell 126, 663–676 (2006). 10.1016/j.cell.2006.07.024

2 Li, H. et al. The Ink4/Arf locus is a barrier for iPS cell reprogramming. Nature 460, 1136–1139 (2009). 10.1038/nature08290

3 Mahmoudi, S. & Brunet, A. Aging and reprogramming: a two-way street. Curr Opin Cell Biol 24, 744–756 (2012). 10.1016/j.ceb.2012.10.004

4 Prigione, A. et al. Mitochondrial-associated cell death mechanisms are reset to an embryonic-like state in aged donor-derived iPS cells harboring chromosomal aberrations. PLoS One 6, e27352 (2011). 10.1371/journal.pone.0027352

5 Lapasset, L. et al. Rejuvenating senescent and centenarian human cells by reprogramming through the pluripotent state. Genes Dev 25, 2248–2253 (2011). 10.1101/gad.173922.111

6 Abad, M. et al. Reprogramming in vivo produces teratomas and iPS cells with totipotency features. Nature 502, 340–345 (2013). 10.1038/nature12586

7 Macip, C. C. et al. Gene Therapy-Mediated Partial Reprogramming Extends Lifespan and Reverses Age-Related Changes in Aged Mice. Cell Reprogram 26, 24–32 (2024). 10.1089/cell.2023.0072

8 Alle, Q. et al. A single short reprogramming early in life initiates and propagates an epigenetically related mechanism improving fitness and promoting an increased healthy lifespan. Aging Cell 21, e13714 (2022). 10.1111/acel.13714

9 Browder, K. C. et al. In vivo partial reprogramming alters age-associated molecular changes during physiological aging in mice. Nat Aging 2, 243–253 (2022). 10.1038/s43587-022-00183-2

10 Ocampo, A. et al. In Vivo Amelioration of Age-Associated Hallmarks by Partial Reprogramming. Cell 167, 1719–1733.e1712 (2016). 10.1016/j.cell.2016.11.052

11 Guan, J. et al. Chemical reprogramming of human somatic cells to pluripotent stem cells. Nature 605, 325–331 (2022). 10.1038/s41586-022-04593-5

12 Hou, P. et al. Pluripotent stem cells induced from mouse somatic cells by small-molecule compounds. Science 341, 651–654 (2013). 10.1126/science.1239278

13 Schoenfeldt, L. et al. Chemical reprogramming ameliorates cellular hallmarks of aging and extends lifespan. EMBO Mol Med 17, 2071–2094 (2025). 10.1038/s44321-025-00265-9

14 Mitchell, W. et al. Multi-omics characterization of partial chemical reprogramming reveals evidence of cell rejuvenation. Elife 12 (2024). 10.7554/eLife.90579

15 Yang, J. H. et al. Chemically induced reprogramming to reverse cellular aging. Aging (Albany NY) 15, 5966–5989 (2023). 10.18632/aging.204896

16 Yücel, A. D. & Gladyshev, V. N. The long and winding road of reprogramming-induced rejuvenation. Nat Commun 15, 1941 (2024). 10.1038/s41467-024-46020-5

17 Mahmoudi, S., Xu, L. & Brunet, A. Turning back time with emerging rejuvenation strategies. Nat Cell Biol 21, 32–43 (2019). 10.1038/s41556-018-0206-0

18 Gorgoulis, V. et al. Cellular Senescence: Defining a Path Forward. Cell 179, 813–827 (2019). 10.1016/j.cell.2019.10.005

19 Chaib, S., Tchkonia, T. & Kirkland, J. L. Cellular senescence and senolytics: the path to the clinic. Nat Med 28, 1556–1568 (2022). 10.1038/s41591-022-01923-y

20 Baker, D. J. et al. Naturally occurring p16(Ink4a)-positive cells shorten healthy lifespan. Nature 530, 184–189 (2016). 10.1038/nature16932

21 Banito, A. et al. Senescence impairs successful reprogramming to pluripotent stem cells. Genes Dev 23, 2134–2139 (2009). 10.1101/gad.1811609

22 Mosteiro, L. et al. Tissue damage and senescence provide critical signals for cellular reprogramming in vivo. Science 354 (2016). 10.1126/science.aaf4445

23 Chiche, A. et al. Injury-Induced Senescence Enables In Vivo Reprogramming in Skeletal Muscle. Cell Stem Cell 20, 407–414.e404 (2017). 10.1016/j.stem.2016.11.020

24 Mosteiro, L., Pantoja, C., de Martino, A. & Serrano, M. Senescence promotes in vivo reprogramming through p16. Aging Cell 17 (2018). 10.1111/acel.12711

25 Ferreirós, A. et al. Context-Dependent Impact of RAS Oncogene Expression on Cellular Reprogramming to Pluripotency. Stem Cell Reports 12, 1099–1112 (2019). 10.1016/j.stemcr.2019.04.006

26 Ogrodnik, M. et al. Guidelines for minimal information on cellular senescence experimentation in vivo. Cell 187, 4150–4175 (2024). 10.1016/j.cell.2024.05.059

27 Heckenbach, I. et al. Nuclear morphology is a deep learning biomarker of cellular senescence. Nat Aging 2, 742–755 (2022). 10.1038/s43587-022-00263-3

28 Saul, D. et al. A new gene set identifies senescent cells and predicts senescence-associated pathways across tissues. Nat Commun 13, 4827 (2022). 10.1038/s41467-022-32552-1

29 Kubben, N., Brimacombe, K. R., Donegan, M., Li, Z. & Misteli, T. A high-content imaging-based screening pipeline for the systematic identification of anti-progeroid compounds. Methods 96, 46–58 (2016). 10.1016/j.ymeth.2015.08.024

30 Acosta, J. C. et al. A complex secretory program orchestrated by the inflammasome controls paracrine senescence. Nat Cell Biol 15, 978–990 (2013). 10.1038/ncb2784

31 Kang, T. W. et al. Senescence surveillance of pre-malignant hepatocytes limits liver cancer development. Nature 479, 547–551 (2011). 10.1038/nature10599

32 von Joest, M. et al. Amphiregulin mediates non-cell-autonomous effect of senescence on reprogramming. Cell Rep 40, 111074 (2022). 10.1016/j.celrep.2022.111074

33 Correia-Melo, C. et al. Mitochondria are required for pro-ageing features of the senescent phenotype. EMBO J 35, 724–742 (2016). 10.15252/embj.201592862

34 Victorelli, S. et al. Apoptotic stress causes mtDNA release during senescence and drives the SASP. Nature 622, 627–636 (2023). 10.1038/s41586-023-06621-4

35 Suryadevara, V. et al. SenNet recommendations for detecting senescent cells in different tissues. Nat Rev Mol Cell Biol 25, 1001–1023 (2024). 10.1038/s41580-024-00738-8

36 López-Polo, V. et al. Release of mitochondrial dsRNA into the cytosol is a key driver of the inflammatory phenotype of senescent cells. Nat Commun 15, 7378 (2024). 10.1038/s41467-024-51363-0

37 Victorelli, S. et al. Mitochondrial RNA cytosolic leakage drives the SASP. Res Sq (2024). 10.21203/rs.3.rs-4876596/v1

38 Miwa, S., Kashyap, S., Chini, E. & von Zglinicki, T. Mitochondrial dysfunction in cell senescence and aging. J Clin Invest 132 (2022). 10.1172/JCI158447

39 Chapman, J., Fielder, E. & Passos, J. F. Mitochondrial dysfunction and cell senescence: deciphering a complex relationship. FEBS Lett 593, 1566–1579 (2019). 10.1002/1873-3468.13498

40 Brand, M. D. & Nicholls, D. G. Assessing mitochondrial dysfunction in cells. Biochem J 435, 297–312 (2011). 10.1042/BJ20110162

41 Martini, H. & Passos, J. F. Cellular senescence: all roads lead to mitochondria. FEBS J 290, 1186–1202 (2023). 10.1111/febs.16361

42 Zhu, Y. et al. The Achilles’ heel of senescent cells: from transcriptome to senolytic drugs. Aging Cell 14, 644–658 (2015). 10.1111/acel.12344

43 Zhu, Y. et al. Identification of a novel senolytic agent, navitoclax, targeting the Bcl-2 family of anti-apoptotic factors. Aging Cell 15, 428–435 (2016). 10.1111/acel.12445

44 Ryan, J. & Letai, A. BH3 profiling in whole cells by fluorimeter or FACS. Methods 61, 156–164 (2013). 10.1016/j.ymeth.2013.04.006

45 Ryan, J., Montero, J., Rocco, J. & Letai, A. iBH3: simple, fixable BH3 profiling to determine apoptotic priming in primary tissue by flow cytometry. Biol Chem 397, 671–678 (2016). 10.1515/hsz-2016-0107

46 Ni Chonghaile, T., et al. Pretreatment mitochondrial priming correlates with clinical response to cytotoxic chemotherapy. Science 334, 1129–1133 (2011). 10.1126/science.1206727

47 Alcon, C. et al. HRK downregulation and augmented BCL-xL binding to BAK confer apoptotic protection to therapy-induced senescent melanoma cells. Cell Death Differ 32, 646–656 (2025). 10.1038/s41418-024-01417-z

48 Sarosiek, K. A. et al. Developmental Regulation of Mitochondrial Apoptosis by c-Myc Governs Age- and Tissue-Specific Sensitivity to Cancer Therapeutics. Cancer Cell 31, 142–156 (2017). 10.1016/j.ccell.2016.11.011

49 Chondronasiou, D. et al. Multi-omic rejuvenation of naturally aged tissues by a single cycle of transient reprogramming. Aging Cell 21, e13578 (2022). 10.1111/acel.13578

50 Lu, J. Y. et al. Prevalent mesenchymal drift in aging and disease is reversed by partial reprogramming. Cell 188, 5895–5911.e5817 (2025). 10.1016/j.cell.2025.07.031

51 Smit, M. A. & Peeper, D. S. Epithelial-mesenchymal transition and senescence: two cancer-related processes are crossing paths. Aging (Albany NY) 2, 735–741 (2010). 10.18632/aging.100209

52 Ansieau, S. et al. Induction of EMT by twist proteins as a collateral effect of tumor-promoting inactivation of premature senescence. Cancer Cell 14, 79–89 (2008). 10.1016/j.ccr.2008.06.005

53 Sahu, S. K. et al. Targeted partial reprogramming of age-associated cell states improves markers of health in mouse models of aging. Sci Transl Med 16, eadg1777 (2024). 10.1126/scitranslmed.adg1777

54 Ulrich, S., Ricken, R. & Adli, M. Tranylcypromine in mind (Part I): Review of pharmacology. Eur Neuropsychopharmacol 27, 697–713 (2017). 10.1016/j.euroneuro.2017.05.007

55 Manzella, N. et al. Monoamine oxidase-A is a novel driver of stress-induced premature senescence through inhibition of parkin-mediated mitophagy. Aging Cell 17, e12811 (2018). 10.1111/acel.12811

56 Ichida, J. K. et al. A small-molecule inhibitor of tgf-Beta signaling replaces sox2 in reprogramming by inducing nanog. Cell Stem Cell 5, 491–503 (2009). 10.1016/j.stem.2009.09.012

57 Maherali, N. & Hochedlinger, K. Tgfbeta signal inhibition cooperates in the induction of iPSCs and replaces Sox2 and cMyc. Curr Biol 19, 1718–1723 (2009). 10.1016/j.cub.2009.08.025

58 Zheng, Y. C. et al. Irreversible LSD1 Inhibitors: Application of Tranylcypromine and Its Derivatives in Cancer Treatment. Curr Top Med Chem 16, 2179–2188 (2016). 10.2174/1568026616666160216154042

59 Carey, B. W. et al. Reprogramming of murine and human somatic cells using a single polycistronic vector. Proc Natl Acad Sci U S A 106, 157–162 (2009). 10.1073/pnas.0811426106

60 Hockemeyer, D. et al. A drug-inducible system for direct reprogramming of human somatic cells to pluripotency. Cell Stem Cell 3, 346–353 (2008). 10.1016/j.stem.2008.08.014

61 Brambrink, T. et al. Sequential expression of pluripotency markers during direct reprogramming of mouse somatic cells. Cell Stem Cell 2, 151–159 (2008). 10.1016/j.stem.2008.01.004

62 Moldoveanu, T. et al. BID-induced structural changes in BAK promote apoptosis. Nat Struct Mol Biol 20, 589–597 (2013). 10.1038/nsmb.2563

63 Chen, Y. et al. SOAPnuke: a MapReduce acceleration-supported software for integrated quality control and preprocessing of high-throughput sequencing data. Gigascience 7, 1–6 (2018). 10.1093/gigascience/gix120

64 Dobin, A. et al. STAR: ultrafast universal RNA-seq aligner. Bioinformatics 29, 15–21 (2013). 10.1093/bioinformatics/bts635

65 Liao, Y., Smyth, G. K. & Shi, W. featureCounts: an efficient general purpose program for assigning sequence reads to genomic features. Bioinformatics 30, 923–930 (2014). 10.1093/bioinformatics/btt656

66 Liao, Y., Smyth, G. K. & Shi, W. The Subread aligner: fast, accurate and scalable read mapping by seed-and-vote. Nucleic Acids Res 41, e108 (2013). 10.1093/nar/gkt214

67 Love, M. I., Huber, W. & Anders, S. Moderated estimation of fold change and dispersion for RNA-seq data with DESeq2. Genome Biol 15, 550 (2014). 10.1186/s13059-014-0550-8

68 Benjamini, Y. & Hochberg, Y. Controlling the False Discovery Rate: A Practical and Powerful Approach to Multiple Testing. Journal of the Royal Statistical Society. 57, 289–300 (1995). 10.1111/j.2517-6161.1995.tb02031.x

69. Data Analysis , bookTitle= ggplot2: Elegant Graphics for Data Analysis. (Springer International Publishing, 2016).

70 Stephens, M. False discovery rates: a new deal. Biostatistics 18, 275–294 (2017). 10.1093/biostatistics/kxw041

71. DEGreport: Report of DEG analysis. R package version 1.13.8. (2017).

72 Mootha, V. K. et al. PGC-1alpha-responsive genes involved in oxidative phosphorylation are coordinately downregulated in human diabetes. Nat Genet 34, 267–273 (2003). 10.1038/ng1180

73 Subramanian, A. et al. Gene set enrichment analysis: a knowledge-based approach for interpreting genome-wide expression profiles. Proc Natl Acad Sci U S A 102, 15545–15550 (2005). 10.1073/pnas.0506580102

74 Kolberg, L. et al. g:Profiler-interoperable web service for functional enrichment analysis and gene identifier mapping (2023 update). Nucleic Acids Res 51, W207–W212 (2023). 10.1093/nar/gkad347

75 Shannon, P. et al. Cytoscape: a software environment for integrated models of biomolecular interaction networks. Genome Res 13, 2498–2504 (2003). 10.1101/gr.1239303

